# Constitutive activation and allosteric mechanisms underlying Gi/Gq signaling bias in SUCNR1 revealed by AlphaFold-based modeling and enhanced sampling simulations

**DOI:** 10.1101/2025.06.08.658530

**Authors:** Aslihan Shenol, Jacob E. Petersen, Michael Lückmann, Thomas M. Frimurer, J Andrew McCammon, Thue W. Schwartz, Wen Ma

**Affiliations:** Novo Nordisk Foundation Center for Basic Metabolic Research, University of Copenhagen, Copenhagen, Denmark; Department of Chemistry and Biochemistry, and Department of Pharmacology, University of California San Diego, La Jolla, California, United States; Department of Physics and Cellular, Molecular, and Biomedical Sciences Program, University of Vermont, Burlington, Vermont, United States

## Abstract

Succinate Receptor 1 (SUCNR1) is a key metabolite sensing GPCR activated by high local succinate concentrations through Gi and Gq signaling pathways. By combining AlphaFold-based modeling, Gaussian accelerated molecular dynamics simulations (72μs in total), receptor mutagenesis, and signaling assays, we here investigate the spontaneous and G-protein binding-associated activation mechanism of SUCNR1. Our molecular simulations predict that, in the absence of G protein or ligand, human SUCNR1 (hSUCNR1), in contrast to murine SUCNR1 (mSUCNR1), adopts canonical active- like conformations in its intracellular domains and undergoes critical conformational changes in its extracellular domain. This finding is consistent with our signaling-assay results showing that hSUCNR1, unlike mSUCNR1, displays a high degree of constitutive signaling. When complexed with either Gαi or Gαq, both h- and mSUCNR1 adopt highly energetically favorable conformations in the extracellular domain, characterized by the unlocking of extracellular loop 2b (ECL2b) and an expanded ligand entry pathway. Interestingly, the simulations reveal that helix 5 of Gq binds less firmly to the TM3/6 cleft of mSUCNR1, resulting in an unstable extracellular domain of the Gq- mSUCNR1 complex, which however can be stabilized by agonist binding. This result is supported by our BRET assays, which show that, in contrast to hSUCNR1, mSUCNR1 fails to recruit mini-Gq to the cell surface in the absence of an agonist. Furthermore, the signaling effects of key receptor residues at G-protein binding site and within ECL2b, as predicted by our simulations, were confirmed by mutagenesis assays. Overall, our integrative approach demonstrates that hSUCNR1 can spontaneously adopt all key conformations associated with receptor activation, and that G protein binding further primes the extracellular receptor domain for agonist binding, which in turn stabilizes the active complex.

## Introduction

G-protein coupled receptors (GPCRs) are the most abundant family of membrane proteins in the human genome with more than 800 members, which through their seven transmembrane-helical structures undergoing dynamic changes can transmit signals across the cell membrane[1]. With their important role in physiology and disease, GPCRs are attractive drug targets and in fact, are the targets for approximately one-third of all medical drugs[2]. Over the last decade, there has been a growing emphasis on the study of metabolite-sensing GPCRs[3]. Research efforts have validated the critical physiological roles of these receptors, in the modulation of e.g. cancer metabolism and regulation of the immune response [4–6]. The significance of these receptors is further underscored by the rising number of resolved structures for Free Fatty Acid Receptors (FFARs), Hydroxycarboxylic Acid Receptors (HCARs), Purinergic Receptors (P2Ys), and the Succinate Receptor 1 (SUCNR1)[7–12]. Succinic acid is a key metabolite and part of the Tricarboxylic Acid (TCA) cycle but metabolic stress conditions, e.g. hypoxia, succinate accumulates due to the reverse action of succinate dehydrogenase. As a result, it is transported out of the mitochondria and the cells in metabolic stress to be sensed by SUCNR1/GPR91[3]. Initially recognized as part of the purinergic receptor family, GPR91 was reclassified in 2004 by He et al. as a succinate receptor[13]. Through autocrine and paracrine way succinate activates SUCNR1 which is associated with beneficial tissue repair and remodelling [14–17]. However, when succinate remains elevated locally, it can lead to harmful inflammatory effects via SUCNR1 present on pro-inflammatory M1 macrophages and activated stellate cells[18]. SUCNR1 can couple to and initiate signaling via both Gi and Gq signaling pathways[19]. It has been shown that the immunomodulatory function of SUCNR1 through hyperpolarization of the M2 macrophages is mediated through the Gq pathway[20]. Despite attempts to develop pharmacological agonist tools/ drug candidates, these have only been successful in signaling through Gi but not the Gq pathway, developing an antagonist able to inhibit murine and human SUCNR1 has also been challenging[21, 22].

Previously, we had through conventional molecular dynamics (MD) simulations, metadynamics analysis, and mutagenesis elucidated the intricate ligand-binding pathway of succinate directed by a cluster of five arginine residues located around the extracellular pole of TM-6 of SUCNR1 [23]. Hereby we identified two low energy, high affinity binding sites for succinate, one in the extracellular vestibule (ECV) and a relatively deep orthosteric site. The ECV site functions both as an initial catching site for the succinate binding path down to the orthosteric site and as a high- affinity binding site, which allows for two succinate molecules to be bound simultaneously. However, these studies did not allow for the identification of the conformational changes in the receptor structure associated with receptor activation as these occur over a larger timescale beyond the reach of conventional MD simulations[24].

To date there are 3 X-ray crystallography structures available of SUCNR1, one of rat SUCNR1(rSUCNR1) in an apo inactive form (pdb:6ibb)[11] and two are antagonist bound inactive structures of humanised - rat SUCNR1 (pdb: 6rnk and 6z10) [10, 11]. Given the lack of available active-state G-protein-coupled structures of SUCNR1 (in the process of submission of this manuscript - active-state structures of hSUCNR1 were reported[25, 26]) we turned in the present study to model the structure of SUCNR1 G protein complexes by AlphaFold2 and the use of the enhanced sampling methods in particular Gaussian accelerated molecular dynamics (GaMD) simulations[27]. Because the original AlphaFold program [29] can only generate a single snapshot for a given GPCR, we applied a multi-state AlphaFold protocol [28] designed for modeling multiple functional states of GPCR (see Methods). Using a state-annotated GPCR template database, this protocol generates inactive structures and G-protein bound structures that can be used as starting points for subsequent GaMD simulations.

GaMD has previously been used to successfully reveal ligand binding, protein folding, protein conformational changes, and GPCR activation mechanisms in detail [30–34]. The basis of this is the application of a harmonic boost potential which subsequently reduces the free energy barriers, accelerating the simulations by orders of magnitude[34]. The acceleration is achieved by adding the boost to dihedral torsions, the system’s total potential energy, or both. The advantage of this method compared to other accelerated methods is that it does not require choosing collective variables (CVs) which allows for it to be used for simulation without initial knowledge about the conformational changes or processes of the system[27].

As our signal transduction assays reveal that human SUCNR1 displays rather clear constitutive activity as opposed to the murine receptor, we first focus on comparing the spontaneous conformational changes of these two very similar receptors without any G protein or agonist present. Our hypothesis is that according to our preferred allosteric selection model for GPCR activation, the receptor should by itself be able to adopt the active conformation, which then is stabilized by either the G-protein and/or the agonist. Secondly, because SUCNR1 requires higher concentrations of succinate to activate Gq signaling as compared to Gi signaling, we test whether we could observe differences in the free energy landscape of SUCNR1 in complex with Gi and Gq. Furthermore, to verify the key allosteric residues involved in G-protein coupling identified in the simulations, we assess the impact of the corresponding mutations using signaling assays. Lastly, we investigate the effects of succinate binding on the receptor conformational activation.

## Results

### AlphaFold-based modeling provides initial structures for enhanced sampling molecular dynamics

In the following, we investigate the molecular activation mechanisms of human and murine SUCNR1 under several conditions: without any G protein or agonist, in complex with either Gi or Gq, and with succinate docked into the orthosteric binding site.

Firstly, we modelled inactive state structures of human and murine SUCNR1 using Multistate AlphaFold2, followed by minimization and relaxation (**see Methods and Supp.Fig. 1A-C**). The predicted inactive structures are closely aligned with the experimentally determined inactive reference structure, i.e. the X-ray structure of ratSUCNR1 (PDB:6ibb) as shown in **Figure 1B**. Similarly, the AlphaFold predicted active SUCNR1 displayed similar key structural changes as observed in the active form of HCAR2 and ß2 adrenergic receptors. Initially, we investigated the spontaneous transition from the inactive state to the active state of hSUCNR1 and mSUCNR1(and rat SUCNR) through 27 simulations, 21,600 ns (Table I). The key conformational changes associated with receptor activation, which we focus on in the present study are shown in **Fig.1**: The canonical opening of the cleft between the intracellular poles of TM-3 and -6 to accommodate helix5 of the G𝛼 subunit **(Fig. 1A)**; the unlocking of ECL-2, which we previously have observed to be associated with agonist activation of SUCNR1[23] **(Fig. 1A)**; and the rotation of Arg^3.50^ of the DRY and the rotation of Tyr7.53 of the NPxxY motifs bringing these two residues in close proximity **(Fig. 1A)**.

**Figure 1:**
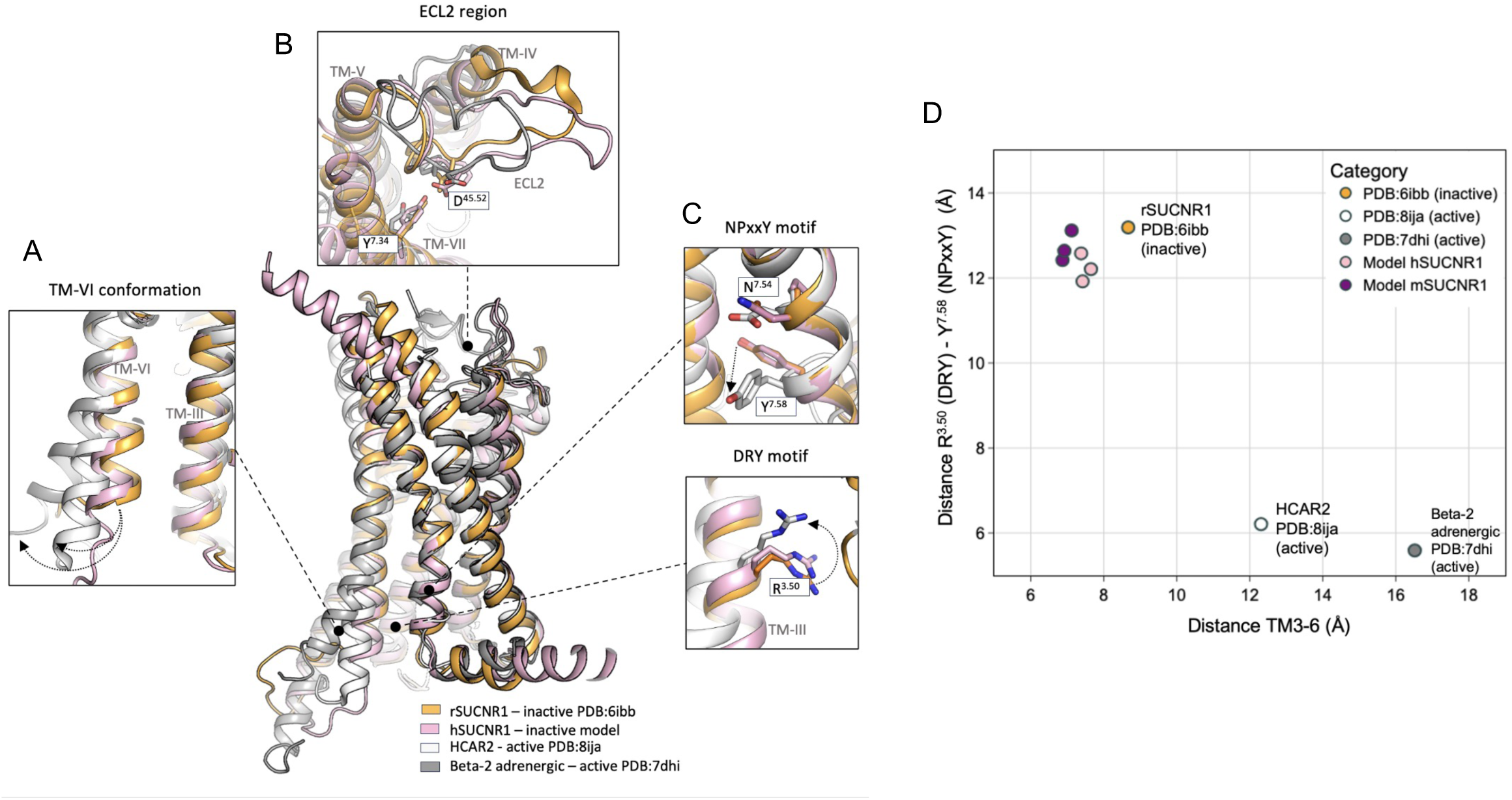
**Modelled Inactive Structures of hSUCNR1 and mSUCNR1 Exhibit Conformational States Similar to Class A GPCRs in the Inactive State** Visual representation of conformational changes related to GPCR activation, showing a comparison between the inactive structure of rSUCNR1 (PDB: 6ibb, orange) to the active structures of HCAR2 (PDB:8ija, white) and beta-2 adrenergic receptor (PDB:7dhi, grey) to the model structure of hSUCNR1 in inactive form (pink). Important conformational motifs are visualized in the panels **A:** TM-6 conformation **B:** ECL2 conformation **C:** NPxxY and DRY motifs. **D:** Modelled hSUCNR1 and mSUCNR1 structures cluster around the reference rSUCNR1 in terms of their TM3-6 distance and DRY-NPxxY motifs’ conformation.

GaMD simulations were subsequently performed with human and murine SUCNR1 in complex with Gαi or Gαq proteins through 44 simulations, totalling 35,200 ns (Table II) to elucidate G protein-associated activation mechanisms of SUCNR1 and aiming at identifying hotspots of interactions that affect the receptor signaling pathways. Lastly, we simulated receptor-G protein complexes with either Gαq or Gαi plus succinate bound in the orthosteric binding pocket across 20 simulations, totalling 16,000 ns (Table III). During all simulations for G-protein coupled systems, the G-protein remained bound to the receptor.

### In contrast to mSUCNR1, the apo form of hSUCNR1 exhibits spontaneous activation, characterized by the opening of its intracellular helices

The structurally highly similar human and murine SUCNR1 signal through both Gi and Gq. Both hSUCNR1 and mSUCNR1 display spontaneous (constitutive) signaling. Importantly, the constitutive signaling of hSUCNR1 is signifcantly greater than that of mSUCNR1 in both Gq signaling (∼10% vs. 4 %) and Gi signaling (∼40% vs. ∼25%) (**Fig. 2A**). Interestingly, when stimulated with succinate, mSUCNR1 is more potent and efficacious than hSUCNR1 in Gi signaling, while the Gq response is comparable between the two species (**Supp.** Fig 1D). Therefore, we would like to see if GaMD simulations allow us to reveal the mechanisms underlying the differential propensity for spontaneously activation in hSUNR1 vs mSUCNR1.

**Figure 2.**
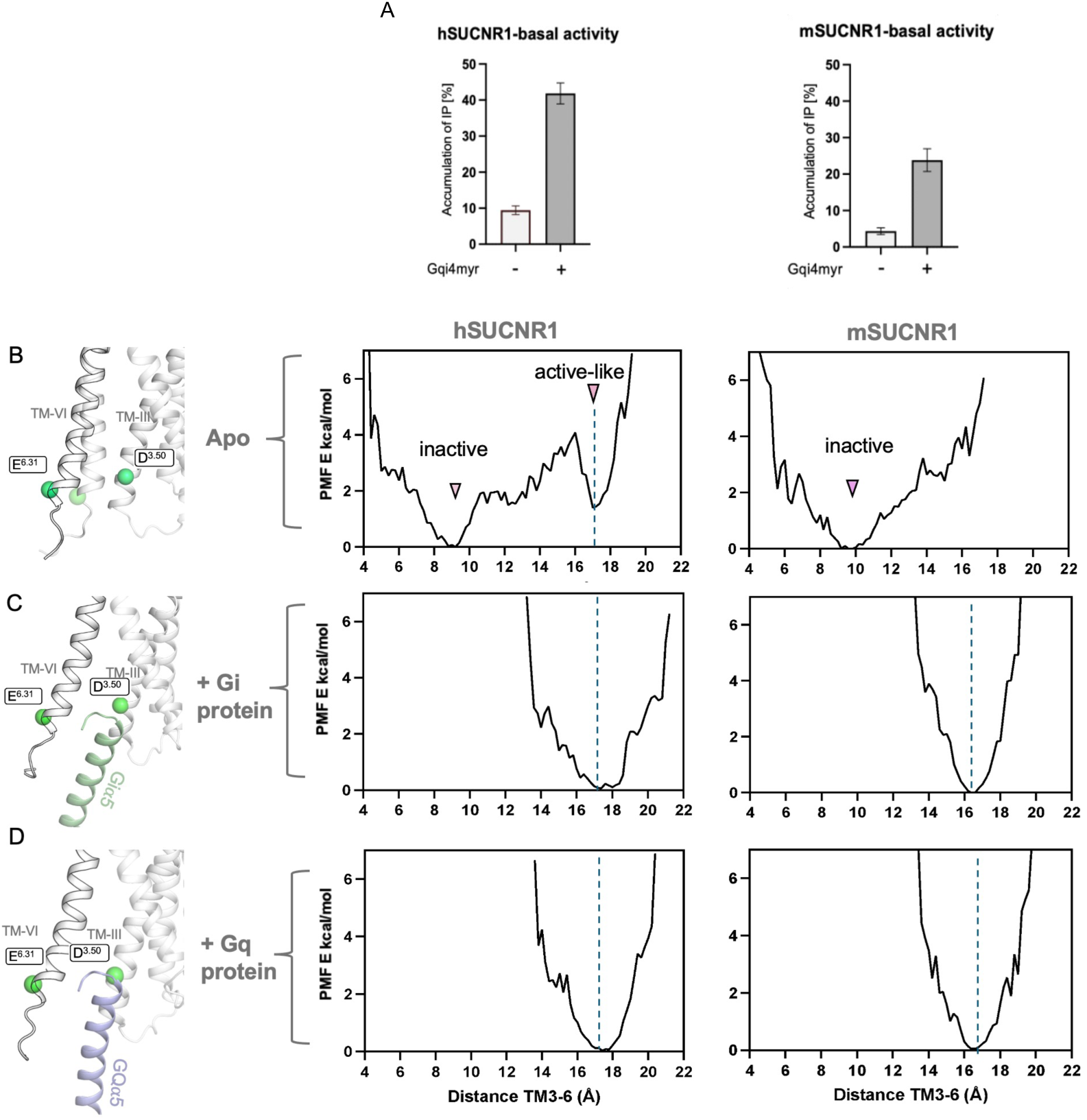
Energy Profiles Predict SUCNR1 Basal Activity, Consistent with Experimental Observations. (A) Basal activity levels of human SUCNR1 (hSUCNR1) and murine SUCNR1 (mSUCNR1) for the Gq (light gray) and Gi (dark gray) signaling pathways. hSUCNR1 demonstrates significant basal activity, while mSUCNR1 shows minimal activity. **(B)** PMF plot depicting the distance between transmembrane helices III and VI for hSUCNR1 and mSUCNR1 in their apo states. hSUCNR1 exhibits two energy minima: ∼9 Å (inactive state) and ∼17 Å (active-like state), as indicated by arrows. In contrast, apo-mSUCNR1 shows a single energy minimum at the inactive state. The inset illustrates the conformational transition of TMVI from the closed (inactive) to open (active-like) state, with the measurement points highlighted. **(C)** PMF plots for the TMIII–VI distance in hSUCNR1 and mSUCNR1 coupled to a Gi protein. Both receptors exhibit energy minima around ∼16.5 Å (mSUCNR1) and ∼17.5 Å (hSUCNR1), consistent with the active-like state observed in apo- hSUCNR1. **(D)** PMF plots for the TMIII–VI distance in hSUCNR1 and mSUCNR1 coupled to a Gq protein. Similar to Gi coupling, energy minima are observed at ∼16.5 Å (mSUCNR1) and ∼17.5 Å (hSUCNR1), aligning with the active-like state of apo-hSUCNR1.

To examine spontaneous activation-like conformational changes of the apo states of the receptor, without ligand and G-protein, we first examined changes in the distance between TM-3 (Cα of R^3.50^) and TM-6 (Cα of E^6.31^) corresponding to the canonical opening of the intracellular cavity during receptor activation. For hSUCNR1 this is observed in 8 of 11 simulations but only happened in 2 of the 11 simulations with mSUCNR1 (**TABLE S1 and** Supp.Fig. 2). This suggests that hSUCNR1 has a higher tendency towards spontaneous active state transition than mSUCNR1. To further validate this, we performed Potential of Mean Force (PMF) analyses across all simulations with hSUCNR1 and mSUCNR1 **(Fig. 2B-D).** First, we measured PMF over the TM3-6 distance for both hSUCNR1 and mSUCNR1 in the apo-state (**Fig. 2B**). The analyses showed that there are two well-defined energy minima (**Fig. 2B - indicated with arrows**), the first one at a TM3-6 distance of ∼9Å corresponding to the inactive state distance between the helices and more importantly, a second minimum at a helical distance of ∼17Å (active-like state) showing an energetically favoured spontaneously opened intracellular cavity corresponding to an outwards shift of TM-6 of - in this case approx. 8 Å **(Fig. 2B - left panel)**. In contrast, the same analysis for mSUCNR1 demonstrated only a single energy minimum at 9 Å corresponding to the closed state **(Fig. 2B),** indicating that the open state with an outward shift of TM-6 is not energetically favoured for apo-mSUCNR1. Thus, the distance in the inactive apo state for both receptors was ∼9Å which corresponds to the TM3-6 distance in the inactive rat SUCNR1 apo-crystal structure (pdb:6ibb).

We then examine the active conformation the h- and mSUCNR1 when coupled to Gi or Gq alpha subunit **(Fig. 2, C and D)**. The PMF plots demonstrated a single, energy minimum corresponding to a TM3-6 distance of ∼17Å for hSUCNR1 and ∼16Å for mSUCNR1 coupled to either Gi or Gq. Importantly, for hSUCNR1 this corresponds closely to the energetically favoured “active-like” state for the TM3-6 distance spontaneously adopted in the apo-form of the receptor. This demonstrates that the intracellular cavity of hSUCNR1 tends to spontaneously transition to a metastable active-like state even in the absence of G-protein or ligand in contrast to mSUCNR1, which tends to stay in its energetically favoured inactive state, agreeing with the observed higher degree of constitutive signaling observed in hSUCNR1.

### The apo form of hSUCNR1 also displays a spontaneous preference for active conformations in the DRY and NPxxY motif as opposed to mSUCNR1

In the active state of GPCRs, the Arg of the DRY motif in TM3 typically shifts from a downward position—where it interacts with the neighbouring Asp —to an upwards-pointing position, where it engages with the Tyr of the NPxxY motif in TM7 (**Fig. 1 C**).

In the starting conformations, the DRY and NPxxY motifs were in the inactive conformations for both SUCNR1 apo-forms **(Fig. 3 A and B, top panels)**. The simulation data highlight a key initial event in the activation of hSUCNR1, where the distance between the residues R^3.50^ and D^3.49^ is increased and the interaction between them is disrupted distance (**Fig. 3A and Supp.Fig. 3 and S4).** This disruption allows R^3.50^ to shift upwards towards Y^7.53^ (NPxxY motif) and the distance between them to decrease (R120-Y295). This repositioning ultimately results in a newly formed interaction between R^3.50^ and Y^7.53^ and the receptor adopting an active conformation (**Fig. 3A and Fig. S4**). This full rearrangement of R^3.50^ and Y^7.53^ was observed in 4 out of the 11 simulations of the apo-hSUCNR1 simulations, but in none of the similar simulations with apo-mSUCNR1 (**TABLE I**), In apo- mSUCNR1 the interaction between R^3.50^ and D^3.49^ remained intact throughout the simulation, and the distance between R^3.50^ and Y^7.53^ remains relatively constant, indicating no observed progression towards receptor activation **(Fig. 3B and Supp.Fig. 3 and 4).** Further, we investigated the energy landscapes obtained from all simulations in relation to the distances between R^3.50^ and D^3.49^ and Y^7.53^, to compare the conformational shifts in hSUCNR1 and mSUCNR1. For hSUCNR1, the energy landscape revealed three distinct major and one minor energy minima **(Fig. 3C indicated with points)**. These correspond to 1) the inactive state characterised by proximity between R^3.50^ and D^3.49^ and a significant distance to Y^7.53^, which is separated by a low energy barrier from 2) a major, low energy intermediate state (IS1) where R^3.50^ is rotated up (the R^3.50^ to D^3.49^ distance increase from approx. 4 to 6.5 Å) and 3) an energetically shallow intermediate state (IS2), which appear to constitute a low energy barrier to 4) the final, deep low-energy active state where the distance between R^3.50^ and Y^7.53^ has decreased from approx. 13Å to 6 Å (**Fig. 3C**). In contrast, the energy landscape of mSUCNR1 shows only a single dominant energy minimum that coincides with the receptor’s inactive conformation with only minimal tendency towards breakage of the R^3.50^-D^3.49^ interaction **(Fig. 3D).** This again indicates a lack of spontaneous dynamic conformational transition towards activation in these motifs in the murine SUCNR1 as opposed to the human receptor.

**Figure 3:**
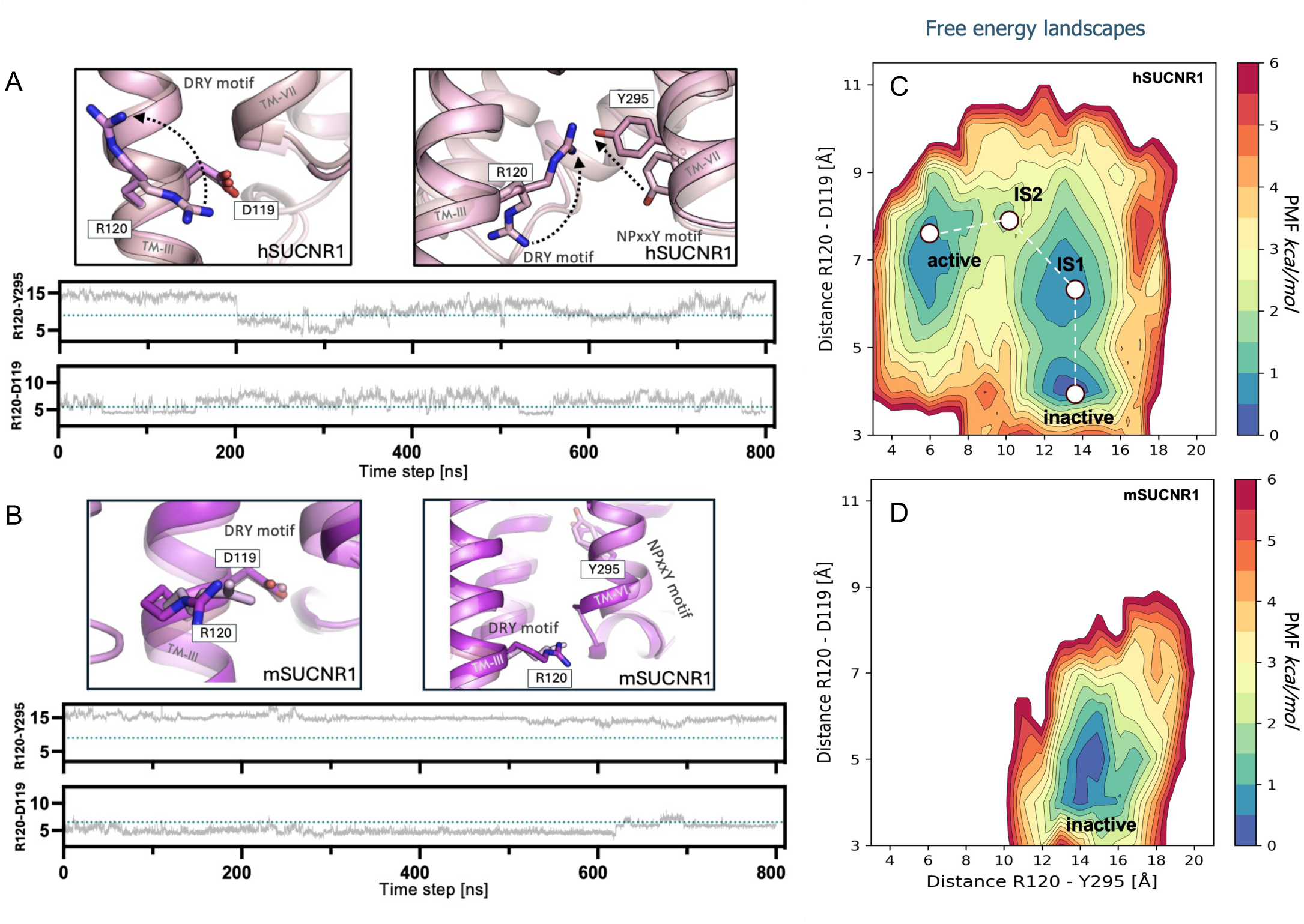
DRY and NPxxY Motif Dynamics Define Receptor Activation Propensity in hSUCNR1 and mSUCNR1. (A) Distance changes between R^3.50^-D^3.49^ (DRY motif) and R^3.50^-Y^7.58^ (DRY to NPxxY) during a representative human SUCNR1 simulation. Initial R3.50-D3.49 interaction weakens while R^3.50^-Y^7.58^ interaction strengthens, indicating conformational shifts toward activation (top panels). **(B)** Murine SUCNR1 shows stable R^3.50^-D^3.49^ interaction and consistently large R^3.50^-Y^7.58^ distance, maintaining an inactive state (top panels). **(C)** Energy landscape for hSUCNR1, based on cumulative simulation data, reveals four energy minima representing inactive, intermediate (IS1, IS2), and active conformations, characterized by changes in R^3.50^-D^3.49^ and R^3.50^-Y^7.58^ distances. **(D)** mSUCNR1 energy landscape, also based on cumulative simulation data, highlights a single energy minimum corresponding to the inactive state, with short R^3.50^-D^3.49^ and large R^3.50^-Y^7.58^ distances.

**Figure 4.**
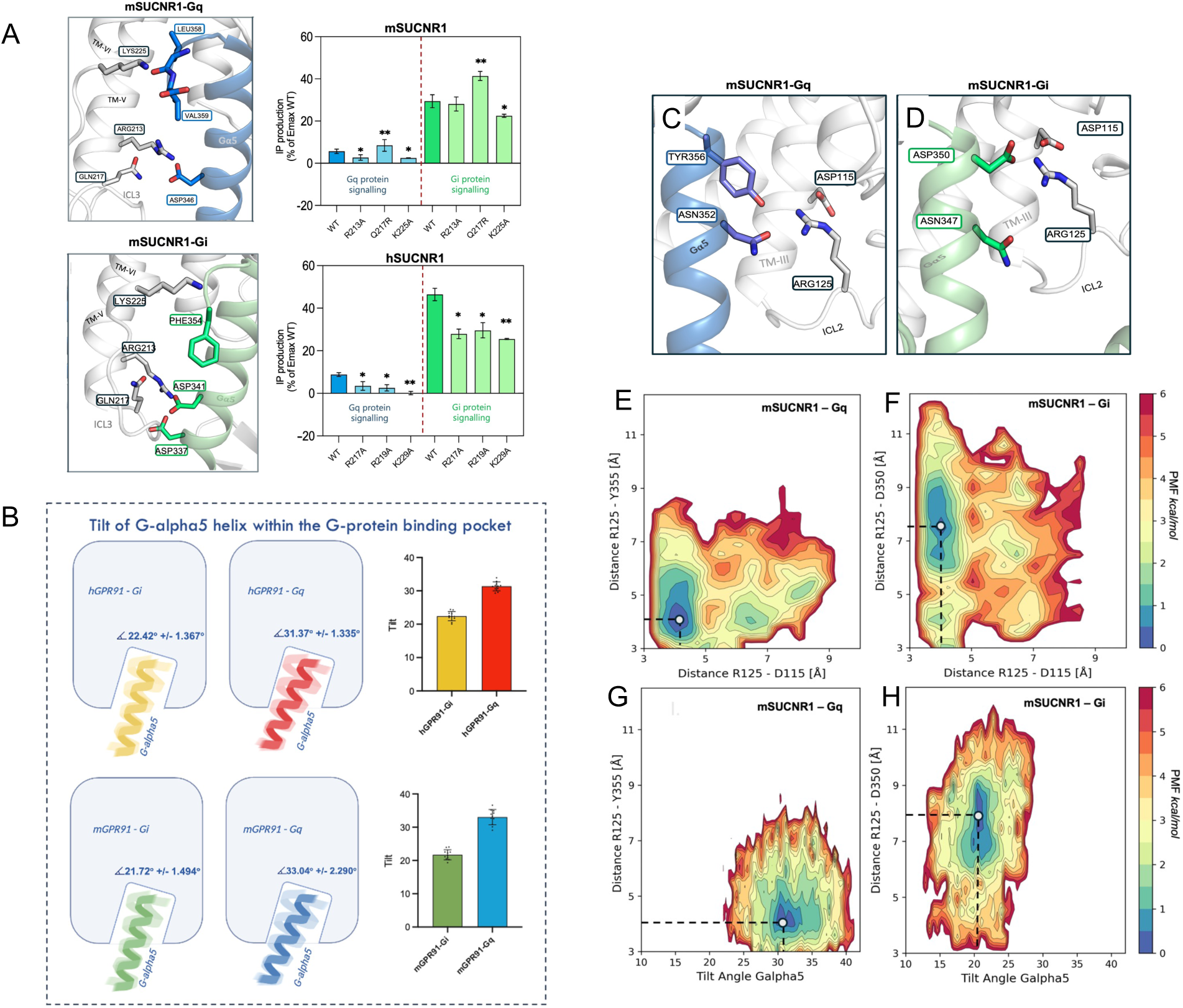
Distinct Dynamics of the G Protein α5 Helix in the Intracellular Cavity Drive Differential Signaling of hSUCNR1 and mSUCNR1. (A) Left panels: Visual representations of the intracellular part of hSUCNR1 and contacts formed between the receptor and the α5 helix of Gq and Gi proteins. Right panels: Bar charts showing the effects of mutations in the intracellular region of hSUCNR1 and mSUCNR1 on basal activity for both Gi and Gq signaling pathways. Changes in basal activity are presented relative to wild-type receptors. Error bars represent SD. **(B)** Comparative analysis of the tilt angle of the G protein α5 helix in hSUCNR1 for Gi (yellow) and Gq (red) proteins, and mSUCNR1 for Gi (green) and Gq (blue) proteins. The results reveal that mSUCNR1 coupled to Gq displays a greater tilt and increased fluctuations compared to Gi coupling. A schematic and bar graph highlight a significantly larger tilt angle for Gq. Error bars represent SD. **(C-D)** Visual representations of interactions between R125 (ICL2) and the α5 helix for Gi and Gq proteins in hSUCNR1 and mSUCNR1, respectively. **(E-H)** PMF plots illustrating the relationship between the distances of R125^34.52^-D115^3.49^ (DRY motif) and R125^34.52^-D350/Y355. For mSUCNR1-Gi **(E),** the favoured conformation includes a salt bridge between R125 and D115 (∼4 Å) and a larger R125-D350 distance (∼8 Å). For mSUCNR1-Gq **(F)**, a similar salt bridge forms between R125 and D115, along with an additional hydrogen bond between R125 and Y355 (∼4 Å), stabilizing Gq coupling. **(G)** PMF plot of the R125^34.52^-D350 distance relative to the α5 helix tilt angle for mSUCNR1-Gi. The favored state shows a tilt angle of ∼21° without R125-D350 interaction (∼8 Å). **(H)** PMF plot of the R125-Y355 distance relative to the α5 helix tilt angle for mSUCNR1-Gq. The energetically favored state includes a tilt angle of ∼32° with an R125-Y355 interaction, stabilizing the Gq-specific α5 helix tilt

### Enhanced Gi vs. Gq binding stability to SUCNR1 and identification of epitopes responsible for the high constitutive signaling of the human receptor

At the time of these analyses, there were no resolved structures of human or murine SUCNR1 coupled to any G-protein, leaving the specifics of these interactions unknown. To address this gap, we conducted a detailed investigation into the dynamics and interactions between SUCNR1 and the G-proteins Gi and Gq, focusing on key structural elements and their roles in coupling specificity.

We first analyzed the interactions formed between SUCNR1 and the G-proteins, concentrating on the intracellular loop 3 (ICL3), TM6, and the Gα5 helix at the G-protein’s C-terminal end (hook region). Notably, we identified a cluster of positively charged residues on the receptor’s intracellular surface, which appeared to play a crucial role in facilitating G-protein coupling **(Fig. 4A-B, left panels).** Mutational analysis revealed that specific substitutions significantly disrupted these interactions. For hSUCNR1, mutations R217^ICL3^A, R219^ICL3^A, and K229^6.32^A reduced constitutive activity for both Gi and Gq pathways, as evidenced by experimental and computational analyses. Similarly, the equivalent mutations in mSUCNR1, R213^ICL3^A and K225^6.32^A, led to a comparable reduction in activity **(Fig. 4A-B, right panels)**. Interestingly, a single mutation in mSUCNR1, Q217^ICL3^R, mimicking the sequence of hSUCNR1 **(Supp.Fig. 6A)**, resulted in increased basal activity for both pathways, aligning with the observed coupling efficiency of hSUCNR1 in the absence of an agonist **(Fig. 4B, right panels)**.

**Figure 5:**
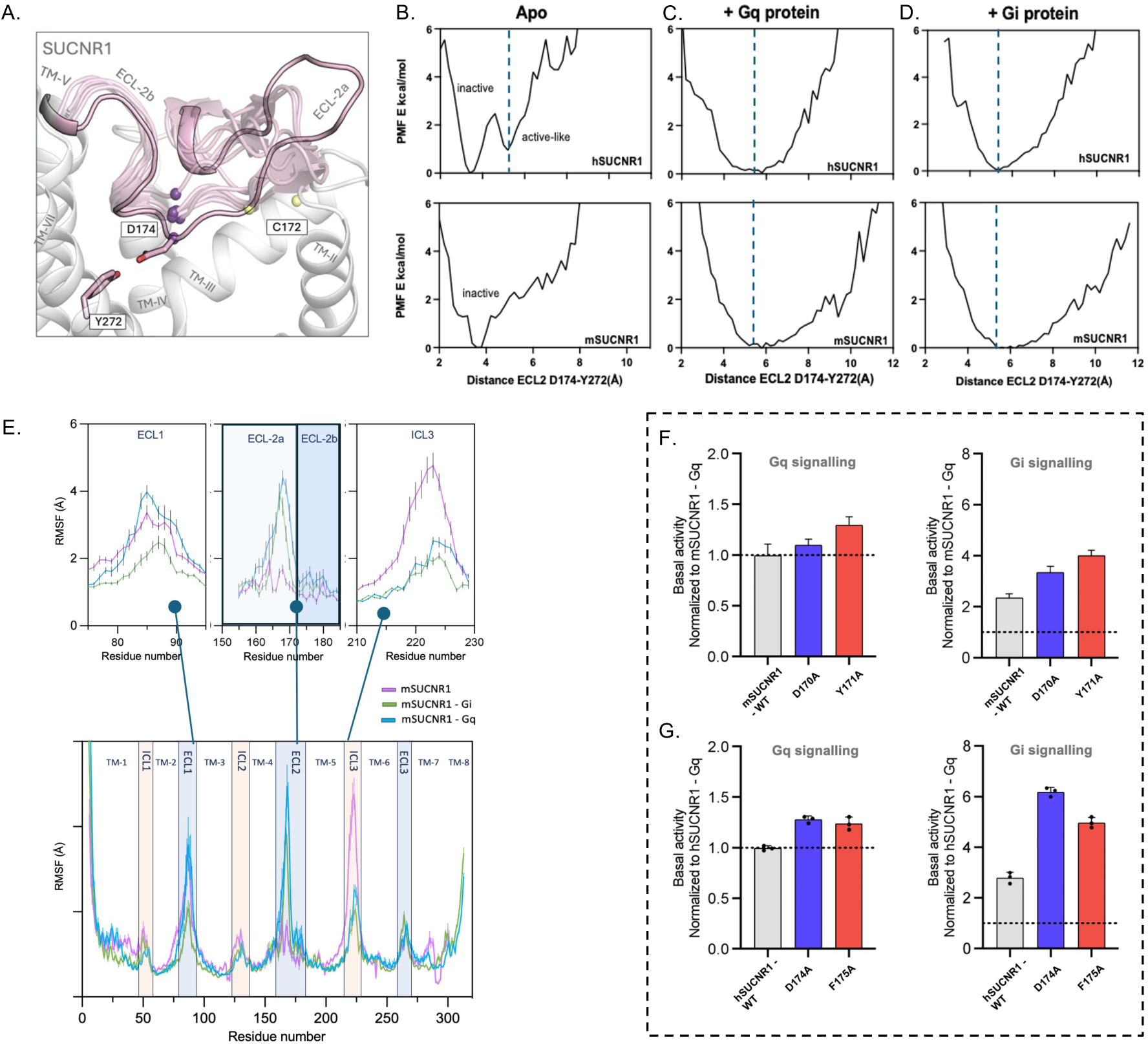
Comparative Analysis of ECL2 Flexibility and the Mutation Effects of Key ECL2 Residues on SUCNR1 Activation. (A) ECL2 transitions from a closed (inactive) to an open (active) state upon activation. Pink: ECL2; purple sphere: D^45.52^ Cα atom; yellow: disulfide bond. **(B)** PMF plots illustrating the distance between Y272^7.35^ and D174^45.52^ for apo hSUCNR1 show energy minima at ∼3 Å (inactive) and ∼5.5 Å (active), suggesting spontaneous ECL2 opening. mSUCNR1 shows a single minimum at ∼3 Å. **(C-D)** With Gi or Gq proteins, both hSUCNR1 and mSUCNR1 favour the open ECL2 state. **(E)** RMSF plots comparing the apo state of mSUCNR1(purple) with Gi- (green) and Gq-coupled systems (blue). ECL1 shows increased flexibility in Gq-protein-bound states and decreased in Gi-bound. while ECL2a fluctuations are significantly higher in G-protein-bound systems and more pronounced in mSUCNR1 than in hSUCNR1 (Supp. Fig). ICL3 flexibility decreases in G-protein-bound mSUCNR1. **F-G:** Effect of single-point mutations D45.52A (blue) and Y/F45.53A (red) on the basal activity of hSUCNR1 and mSUCNR1 in Gi and Gq signaling. Both mutations result in a significant increase in basal activity compared to wild-type levels.

**Figure 6:**
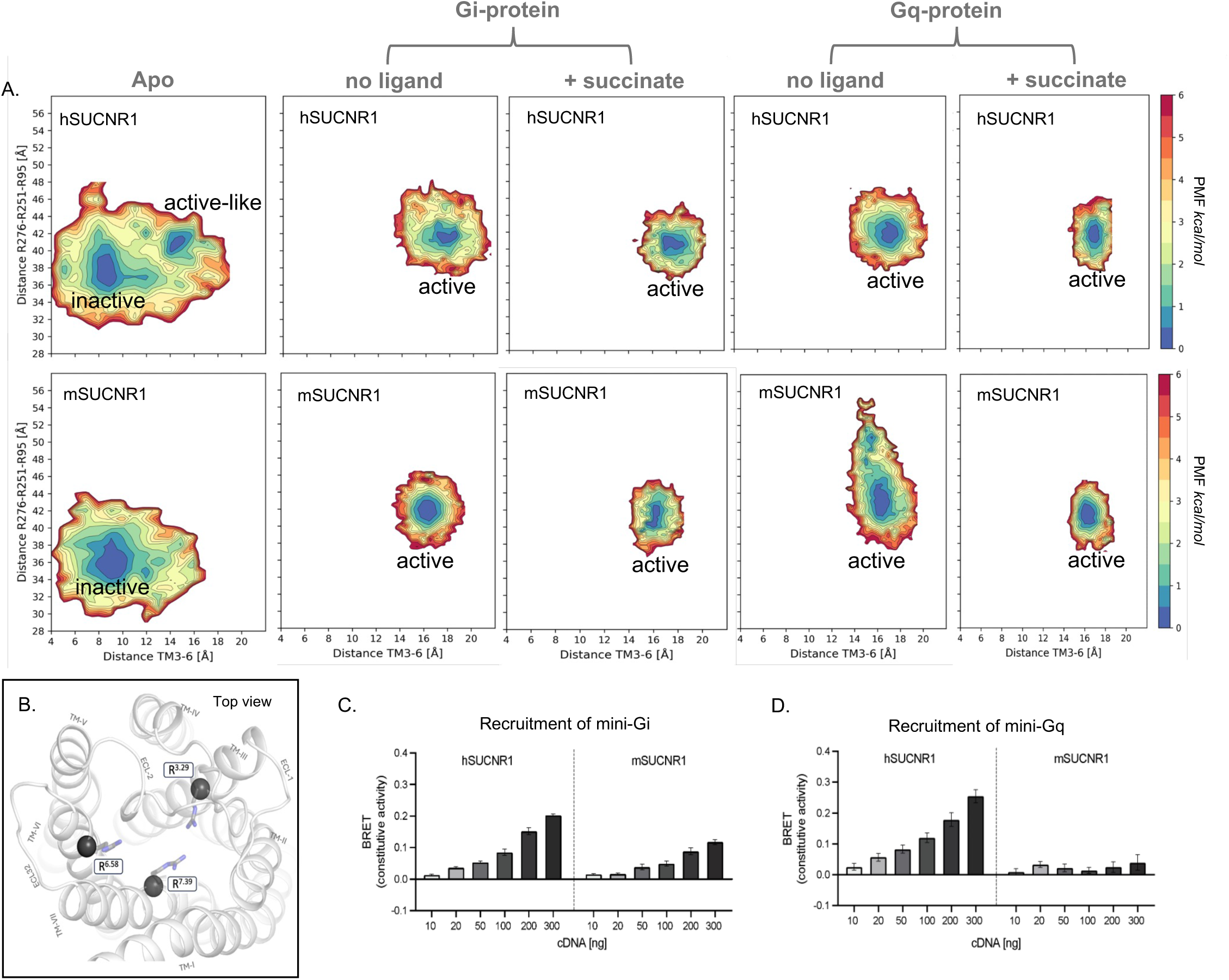
Orthosteric Binding Pocket Dynamics upon G-protein and Succinate Binding Highlight the Allosteric Mechanism of Receptor Activation. A: PMF plots illustrating the energy landscape as a function of the TM3-TM6 distance and triangulation distance (R^3.29^-R^6.58^-R^7.39^) for hSUCNR1 and mSUCNR1 across five systems: Apo, Gi-coupled (± ligand), and Gq-coupled (± ligand).**Apo:** hSUCNR1 favors an expanded orthosteric pocket, while mSUCNR1 prefers a non-expanded configuration. **Gi-coupled (± ligand):** Active state is defined by ∼42Å triangulation distance. Ligand binding sharpens the energy minimum and stabilizes the receptor for both species. **Gq-coupled (without ligand):** hSUCNR1 resembles its Gi-coupled state (∼42Å), whereas mSUCNR1 shows a broader range (40- 52Å). **Gq-coupled (with ligand):** Ligand binding stabilizes mSUCNR1, restoring its energy landscape to resemble the Gi system. **B:** Visualization of SUCNR1’s orthosteric pocket with R^3.29^, R^6.55^, and R^7.39^ shown as sticks, and their Cα atoms as grey spheres. **C-D:** Recruitment of mini-Gi and mini-Gq at the plasma membrane with increasing receptor DNA doses. hSUCNR1 shows enhanced constitutive activity with both G-proteins, while mSUCNR1-Gq exhibits low basal activity with no significant change. Data are shown as mean ± SEM.

To probe the influence of G-protein dynamics on coupling and receptor bias, we performed a comparative analysis of the Gα5 helix within the intracellular cavity of SUCNR1. This included assessing the tilt angle, orientation, and flexibility of the Gα5 helix in both Gi- and Gq-coupled systems. The helix’s tilt angle was determined by measuring the angle formed between the principal axis of the helical bundle of the GPCR and the vector passing through the last ten helical turns of the Galpha5 helix (C-terminal end) as described in [35]**(Fig. 4C).** In the hSUCNR1-Gi complex, the tilt angle of the Gα5 helix was approximately 22° with a standard deviation of ±1.36°, whereas in the hSUCNR1-Gq complex, the angle increased to ∼31° with a SD of ±1.33° **(Fig. 4C, top panels)**. The same trend was observed for mSUCNR1, where Gi complexes displayed a smaller tilt angle and Gq complexes exhibited greater flexibility and a larger tilt angle. These differences were further supported by root-mean-square deviation (RMSD) analyses, which showed higher deviations in the Gα5 helix of Gq-coupled systems compared to Gi **(Supp.** Fig. 6C).Interestingly, quantification of the average tilt angle for both systems across all simulations in the absence of a ligand showed that the Gq-coupled mSUCNR1 systems exhibited a more loosely packed binding pocket, as indicated by the greater flexibility of the Gα5 helix within the receptor’s intracellular cavity **(Fig. 4C, bottom bar chart)**. These findings suggest that the distinct dynamics of the Gα5 helix contribute to the differences in coupling efficiency and receptor preference for Gi and Gq proteins.

To further elucidate the molecular basis of coupling specificity, we analysed residue-specific interactions within the receptor-G-protein complexes which define the tilt angle of coupling. A key observation was the role of residue R125 in ICL2, which formed a salt bridge with D115^3.49^, stabilizing the conformation of ICL2 across all simulated systems **(Fig. 4D-E)**. Initially, in the modelled systems, R125^ICL2^ is positioned in proximity to ASP350 in the Gα5 helix of the Gi protein and TYR356 in the Gα5 helix of the Gq protein. To see if these interactions are important for the orientation of the G-protein we looked at the energy landscape **(Fig. 4F-I).** Notably, in Gq-bound systems, R125^ICL2^ established energetically favoured additional interactions with TYR356 in the Gα5 helix, forming hydrogen bonds that contributed to the stabilization of the greater tilt angle observed in these complexes. In contrast, in Gi-bound systems, the energetically favoured distance between R125^ICL2^ and ASP350 in the Gα5 helix (∼8 Å) suggested no direct interaction, aligning with the less pronounced tilt angle **(Fig. 4F-G).** PMF analyses of the corresponding tilt angle provided further insights into these interactions. The interaction between R125^ICL2^ and TYR356 was associated with an energy minimum at a residue distance of ∼4 Å and a tilt angle of ∼32°, stabilizing the Gq complex **(Fig. 4H).** On the other hand, the formation of interaction between R125^ICL2^ and the Giα5is not energetically favoured for the orientation of the Gi-protein within the intracellular cavity **(Fig. 4I).**

Our findings reveal structural and mutational determinants underlying SUCNR1’s higher basal activity with Gi compared to Gq. Mutations in ICL3 and TM6 (e.g., R217^ICL3^A, R219^ICL3^A, and K229^6.32^A) significantly reduced basal activity for both pathways, highlighting the importance of charged residues in coupling. The humanizing mutation Q217^ICL3^R in mSUCNR1 increased basal activity, aligning with hSUCNR1’s higher basal levels, underlying the importance of this sequence difference for the receptor basal activity. Gi-coupled systems exhibited tighter packing of the Gα5 helix and more stable interactions, while Gq-coupled systems showed increased flexibility and a looser binding pocket. These results connect structural dynamics, key residues, and receptor-G-protein interactions to SUCNR1’s distinct basal signaling profiles.

### Conformational changes of ECL2 associated with SUCNR1 activation occurs both spontaneously and upon G protein binding

ECL2 is part of the activation processes of many Classes A GPCRs [36–38]. We have previously shown that the binding of endogenous ligands and agonists but not of antagonists to SUCNR1 results in conformational changes in ECL2, i.e. opening of the path to the orthosteric binding pocket[23]. In our current simulations we explore the role of ECL2 in the activation of the GPCR in the absence of an agonist.

In the inactive forms of both hSUCNR1 and mSUCNR1, ECL2 adopts a closed conformation stabilized by an interaction between residues D174^45.52^ and Y277^7.35^ (**Fig. 5A and Supp.** Fig.1 **A**). Simulations revealed that this bond spontaneously breaks, causing the ECL2b segment to shift away from the orthosteric pocket and TM-7 (**Supp.Fig. 5**). PMF analysis showed a clear energy preference for the open state in apo hSUCNR1, whereas mSUCNR1 displayed a tendency to maintain its closed conformation **(Fig. 5B-Apo and Supp.** Figure 5). Furthermore, the Gi or Gq protein-bound forms notably shift the energy minimum to an open ECL2 state for both receptor species, underscoring the role of ECL-2 as an important part of the receptor activation mechanism **(Figure 5B - middle and right panels)**.

To further validate ECL2’s involvement in constitutive activity, we introduced the mutations D^45.52^A and Y/F^45.53^A and measured the basal activity of the receptors in Gi and Gq signaling pathways **(Fig. 5C).** For hSUCNR1-Gi the D^45.52^A mutation increased basal activity twofold compared to wild type, with F^45.53^A showing a significant increase as well. A similar trend was observed for Gq signaling, albeit with a smaller magnitude **(Fig. 5C)**. Interestingly in mSUCNR1, the D^45.52^A increased basal activity compared to wild type but not as much as the same mutation in hSUCNR1. Conversely, the mutation Y^45.53^A, in contrast with the same position in hSUCNR1, increased the basal activity more than the mutation on the aspartic acid, highlighting species-specific differences. We identified the increase of the basal activity of SUCNR1 to be related to the ability of beneficial residue rearrangement in hSUCNR1 which holds the active ECL2 conformation in which i.e. R^6.55^ is free to shift upwards and interact with ECL2 and stabilize the active state of the receptor **(Supp.Fig. 9-12)**, which was not prevalent in mSUCNR1 The Y^45.53^A mutation in mSUCNR1 would allow for this stabilizing rearrangement, resulting in an increased basal activity.

RMSF analyses compared the receptor flexibility of extracellular loops and intracellular regions in apo and G-protein-bound states, across all simulations (**Fig. 5D and Supp.Fig. 7A**). For hSUCNR1 the major RMSF changes were observed mainly in the G-protein coupled systems compared to the apo system. RMSF of ECL1 remained mostly unchanged for all systems with hSUCNR1 **(Supp.Fig. 7A – top left panel).** In contrast, the ECL2a fluctuation significantly increased for hSUCNR1 coupled to any of the G-protein and only a slight increase was recorded for ECL2b **(Supp.Fig. 7A – middle panel).** Finally, the ICL3 fluctuation decreased significantly in G-protein coupled systems **(Supp.Fig. 7A – top right panel)** possibly due to the interactions between the receptor and the protein at that vicinity. Analogues plots for the mSUCNR1 systems showed different overall receptor RMSF compared to its human counterpart **(Fig 5D).** Unlike hSUCNR1 – apo-mSUCNR1 system’s (purple) ECL1 RMSF is significantly elevated, this high fluctuation is retained for mSUCNR1-Gq (blue) but brought down by mSUCNR1-Gi systems (green) **(Fig. 5D – top left panel)**. Comparison of the ECL2 RMSF indicates an increase of ECL2a flexibility for G-protein coupled mSUCNR1 systems to an even greater extent than for hSUCNR1, and a slight increase of the ECL2b flexibility **(Fig. 5D – middle panel).** Similarly to hSUCNR1 ICL3 RMSF decreases significantly in mSUCNR1 coupled to G proteins due to the stabilization of the loop in contact with the G-proteins. **(Fig. 5D – right panel).** The increased fluctuations of most of the extracellular part of mSUCNR1 in the complex with Gq is an indication of a possible destabilization of the receptor structure. These differences can be attributed to sequence divergences in ECL1 and ECL2, the regions with the least sequence conservation between the two species **(Supp.** Fig. 6B). Our findings underscore that ECL2 dynamics are integral to SUCNR1 activation, even in the absence of ligands. The spontaneous opening of ECL2 and its G-protein-induced stabilization highlight its role as a mechanistic switch in receptor activation. Moreover, species-specific sequence differences contribute to variations in ECL2 behaviour and overall receptor-protein complex stability.

### Allosteric regulation of ligand binding pocket by G protein binding revealed by energy landscape analysis and mini-G protein BRET assays

Succinate binding and pathway progression are integral to SUCNR1 activation, involving a series of receptor conformational changes. Previously we have described how succinate initially binds the extracellular vestibule (ECV) and is subsequently translocated to the deep orthosteric site by R251^6.58^ while passing and breaking the constraining H-bond between ECL-2b D170^45.52^ and Y272^7.35^ through an associated water cluster[23]. GaMD has previously been used to describe how G-protein binding can be associated with the expansion of the orthosteric binding site in the M2 muscarinic receptor [39]. In SUCNR1 we focus on the narrow part of the succinate entry path as defined by the triangulation distance between the Cα of R251 and the Cα5 of the two Arg located at the opposite wall or bottom of the orthosteric pocket, R95^3.29^ and R276^7.39^ **(Fig. 6B)**[23]. **Fig. 6A** shows energy landscapes comparing murine and human SUCNR1 across different systems i.e. the apo form and G- protein coupled and now when the focus is on the binding pocket with or without succinate bound in the orthosteric pocket. In the apo systems, the energy landscape revealed that with the spontaneous increase of the intracellular distance TM3-6 **(described in Fig.2)**, the orthosteric pocket located approx. 20 Å above the G-protein binding intracellular cavity, expanded from an inactive to an active state in hSUCNR1 but not in mSUCNR1, (**Fig. 6A – left top panels)**. The introduction of Gαi or -q in both cases favoured an expansion of the ligand binding pocket circumference to ∼42Å, which is only approx. 1Å larger than observed in the spontaneously adopted active state of the hSUCNR1-apo system. However, introducing succinate condensed the energy landscape without altering the pocket circumference. **(Fig. 6A - top panels)**. Conversely, mSUCNR1 displayed notable differences. The introduction of Gαi in mSUCNR1 resulted in an increase in the binding site circumference with an energy minimum of ∼42Å. However, surprisingly the energy landscape for mSUCNR1 in complex with Gαq was very different from that observed with hSUCNR1 as it extended further ‘upward’ towards a binding site circumference between 42 and 50Å,. a state larger than the other active states. Despite the well-defined TM3-6 distance, this larger pocket circumference and corresponding increased ECL **(Fig. 5)** fluctuations indicated destabilization in the mSUCNR1-Gq complex, a phenomenon not observed with Gi coupling. **(Fig. 6A - bottom panels).** Importantly, the introduction of succinate to the mSUCNR1-Gq complex shifted the energy landscape to a similarly condensed picture as observed for succinate bound to the hSUCNR1-Gqcomplex – as expected - stabilizing the orthosteric pocket. Across all simulations, the average triangulation distance was largest for mSUCNR1-Gq and consistent for hSUCNR1-Gi or Gq and mSUCNR1-Gi **(Supp.Fig. 6)** Our analysis suggests that even though mSUCNR1 can form interactions with Gq-protein, the overall structure of the mSUCNR1 complex with Gq is more unstable than the complex with Gi in the absence of a ligand as reflected in the energy landscapes presented in **Fig. 6** and in the RMSF profiles shown in **Fig 5**. Direct measurements of mSUCNR1’s ability to form complexes with Gq and Gi confirmed these findings. Using KRAS as a directional BRET intermediary located at the cell surface, we performed direct measurements of the ability of mSUCNR1 to form complexes with Gq versus Gi by studying recruitment and activation of mini-Gsi and mini-Gsq with mSUCNR1 in comparison with hSUCNR1. In this system we observed strong receptor cDNA dose-dependent constitutive activation for hSUCNR1, demonstrating robust coupling to both mini-Gi and mini-Gq. mSUCNR1 exhibited robust mini-Gi coupling but showed significantly reduced and absent activation with mini-Gq, aligning with the simulation-predicted instability in the mSUCNR1-Gq complex, **(Fig. 6C-D)** demonstrating that the molecular simulations successfully predicted the disparity in constitutive signaling capability between hSUCNR1 and mSUCNR1.

While hSUCNR1 maintains structural stability in G-protein-bound states, mSUCNR1 exhibits Gq-specific destabilization, as evidenced by its energy landscapes, orthosteric pocket dynamics, and reduced constitutive activity. This suggests that sequence differences, particularly in ECL regions, influence species-specific G-protein coupling and receptor activation.

## Discussion

Through AlphaFold modelling, GaMD simulations and signal transduction experimental validations, of both human and murine SUCNR1 we have determined the structural and dynamic basis for 1) ligand-independent, constitutive receptor activation 2) the allosteric mechanism connecting G-protein binding and conformational activation of the extracellular receptor domains, and 3) Gi vs. Gq signaling bias of the receptor.

All GPCRs display some degree of constitutive activity, i.e. they are able to signal in the absence of agonist. This can be a few percent or up to more than 50% constitutive signaling as observed for e.g. the ghrelin or the CB1 cannabinoid receptors [40, 41]. At molecular level, this means that the receptors should be able to undergo the conformational changes from inactive to active states in the absence of an agonist. Experimentally resolved receptor structures do not provide relevant insights into the receptor activation or mechanisms underlying coupling bias because they do not account for the essential receptor flexibility. However, the dynamic nature of the receptors is fundamental to their ability to transmit signals across the cell membrane. Our GaMD findings support the notion that GPCRs, in this case SUCNR1, can transition spontaneously between states in the absence of bound ligand or G-protein. In our simulations, both the intra- and extracellular domains of SUCNR1 spontaneously transition from inactive to active states through multiple intermediate metastable states. Notably, the greater propensity for hSUCNR1 to adopt active conformations spontaneously, compared to mSUCNR1, provides the molecular explanation for its higher basal signaling activity.

The most pronounced conformational changes in GPCR activation occur on the intracellular side and specifically in the G-protein coupling interface. Thus, in the apo-form of the constitutively active human SUCNR1 we observe an energetically favoured, ∼8 Å spontaneous opening of the cleft between TM3 and TM6. This opening corresponds to the gap required to encompass H-5 of the G proteins and is not observed in the murine receptor, which has very low constitutive activity. This conformational change between TM3 and TM6 is as expected associated with a significant structural rearrangements in the conserved DRY and NPxxY motifs and their interactions. It is important to note that these changes are not necessarily observed simultaneously[42]. Often the receptor structure explores different intermediate states until a fully active state is thermodynamically stabilized, supporting the notion of GPCRs existing in a dynamic equilibrium between many different states. In our simulations SUCNR1 goes through multiple metastable states in which one or more of the conformational changes may occur until a stable receptor active state is achieved. This has also been observed in molecular dynamics studies performed on muscarinic[43, 44] and adrenergic receptors[45]. In our case capturing metastable receptor activation states helps us distinguishes the basal activity differences between two species of the same receptor. Additionally, it allowed for deeper understanding of the thermodynamic basis for the residue rearrangement in and between the NPxxY and DRY motifs as displayed in figure 3.

During activation of GPCRs important conformational changes may also occur in the extracellular domains[36]. Our GAMD analysis of SUCNR1 underscores the importance of conformational changes in both ECL2 and the orthosteric agonist binding site and shows how these changes are associated with the intracellular conformational changes and occur both spontaneously and upon binding of the G protein. For SUCNR1 we previously demonstrated that the H-bond lock between Y277 and D174^ECL2^ opens upon succinate and CES binding but not when the antagonist binds in the orthosteric site[23]. Our study demonstrates a spontaneous unlocking of ECL2 through opening of this H-bond in the human (an event not observed in murine apo-hSUCNR1), and that binding of G proteins at the intracellular side stabilizes a fully open loop state in both mSUCNR1 and hSUCNR1. Two residues within ECL2 are identified as being significant for these conformational change, i.e.Asp^45.52^ and Phe/Tyr ^45.53^ Our experimental results, which show that alanine mutations of these residues in ECL2 increased the basal activity of SUCNR1, i.e. its propensity to associate with the G proteins, underlines the two-way allosteric connection between the G-protein binding site and the agonist binding site, importantly even in the absence of an agonist ligand. The observed unlocking of the constrained ECL2 and widening of the orthosteric binding pocket at the extracellular receptor domain upon G protein binding at the intracellular receptor domain aligns well with the established notion that G protein binding in general increases the receptor affinity for agonists.

During the preparation of this manuscript, cryo-EM structures of hSUCNR1 bound to succinate and coupled to Gi and mini Gs/q were published[25, 26]. In these active structures, ECL2 was observed in an unlocked, open state, as opposed to the inactive state structure of SUCNR1[11], confirming our computational and experimental results showing the importance of ECL2 for the activation of the receptor. Similarly, a difference between ECL2 conformation in receptor inactive and active state has been observed in the HCAR2 receptor in a series of recently published structures[12] and in the beta-adrenergic receptors where the loop opening and rearrangement of residues in the corresponding positions were observed upon binding of agonists [38, 46].

Concerning G protein binding, MD simulations by Sandhu and coworkers have previously showed that the C terminal helix of Gs, Gi, and Gq adopt different angular orientations within the receptor intracellular cavity [35]. Our findings regarding the tilt angle of the alpha5 helix in the G protein provide further insights into the coupling specificity or bias of SUCNR1. The tilt angle for the hSUCNR1-Gi systems was around 22°, whereas in complex with Gq it was approx. 31°. This difference suggests that the structural arrangement of the alpha5 helix contributes to G protein coupling efficiency. Crucially, the presence of specific interactions, such as the contact between Arg125^34.53^ in ICL2 of SUCNR1 and Tyr356 in the Gq α5 helix, was identified as a determinant for greater Gq specific contact angle. The predicted difference in the tilt angle of both G-proteins agrees with the results demonstrated by the recently resolved structures [25].

Agonist binding, while not essential for the conformational transition from inactive to active state of GPCRs, modulates their rate and thermodynamic stability. An agonist bound to the orthosteric pocket stabilizes an intracellular active conformation or increases the probability of the intracellular cavity being transformed into an active conformation, to which the receptor can adopt spontaneously. Our analyses revealed that succinate binding stabilizes the active conformation of SUCNR1, which was particularly evident in the expansion of the orthosteric binding pocket for the G protein-coupled states. We observed that binding of G-protein to SUCNR1 induced conformational changes in the orthosteric agonist binding path and pocket, i.e. expanding it to “prepare” for ligand binding, similar to prior MD results reported for muscarinic, beta-adrenergic[47] and canabinoid[48] receptors. Interestingly, in our simulations with mSUCNR1 in complex with Gq, we noted an extremity in the pocket expansion together with increased RMSF values for both ECL2 and ECL1. The data suggested that in complex with Gq (as opposed to Gi), the extracellular part of the receptor is destabilized, which however becomes stabilized upon succinate binding to the orthosteric pocket. This led us to propose that mSUCNR1 likely cannot efficiently couple to Gq in the absence of an agonist to stabilize this conformation. We validated this notion experimentally by showing that in the absence of succinate, mSUCNR1 (as opposed to hSUCNR1) was not able to recruit mini-Gq to the plasma membrane, while it efficiently recruited mini-Gi.

Collectively, our study provides a molecular dynamics perspective on constitutive, ligand independent GPCR signaling and G protein bias. The equilibrium between active and inactive states is pivotal for the receptor’s readiness to respond to both agonist and G protein binding, which in turn biases the equilibrium toward activation and signaling. These findings not only advance our knowledge of GPCR research but also offer valuable information for in silico drug discovery targeting SUCNR1. Furthermore, they underscore the effectiveness of combining AlphaFold-based modeling with enhanced sampling MD simulations to predict activation mechanisms, coupling complexities, and signaling bias in GPCRs.

## Methods

### Computational Methods

#### Model building and refinement

The human and murine SUCNR1 model structures were generated using the multi-state modelling protocol for AlphaFold2 [28]. The protocol extends AlphaFold2 [29] to model either inactive or active receptor states with high accuracy, using GPCR structural templates annotated by state. For the inactive state, we provided either the hSUCNR1 or mSUCNR1 sequence as input and biased the templated-based modeling toward the inactive conformation. After ranking and validating an inactive state structure, we used the Protein Preparation Wizard workflow (Schrödinger[50]) to minimize the structure and assign residue protonation states for pH=7 using PROPKA. Similarly, we also used the AlphaFold Multimer protocol to prepare hSUCNR1 and mSUCNR1 structure coupled to Gi and Gq protein alpha subunit. After validating that the receptors are in an active state we followed with the Protein Preparation Wizard.

#### System building and simulations

Eleven different systems were prepared for simulations - three in an apo state (mSUCNR1, hSUCNR1, and rSUCNR1 - pdb:6ibb) - 4 with Gi or Gq protein alpha subunit coupled to murine or human SUCNR1 and 4 with a coupled G - protein and succinate prebound in the orthosteric pocket. Succinate was further parameterized using Ligprep and docked into the agonist binding pocket of the prepared SUCNR1 structures using Glide Schrödinger[51]. The correct protonation states for histidines were determined using PROPKA at pH 7, ensuring physiological relevance (Schrödinger). Additionally, disulfide bridges were accurately constructed to be the same as those found in the resolved crystal structure of SUCNR1. The Membrane Builder module of the CHARMM-GUI server was used to prepare simulation input systems [52]. To ensure stability and neutrality, all chain termini were capped with acetyl and methylamide groups.

The positioning of receptors within a palmitoyl-oleoyl-phosphatidyl-choline (POPC) bilayer was guided by alignments from the OPM database [53], establishing an optimal membrane environment. The simulation systems were solvated in a 0.15 M NaCl solution to mimic physiological conditions, neutralized, and equilibrated to a temperature of 310 K with the TIP3P water model employed for solvation. The AMBER ff19SB and AMBER LIPID 17 parameter sets were utilized for receptors and lipids to ensure accurate molecular interactions[54] . The GAFF2 parameters were used for succinate.

The final simulation box consisted of ∼137 000 atoms, 115 POPC in the membrane upper leaflet and 119 in the lower leaflet, 92 Na and 92 Cl ions, the box dimensions are approximately 98x150x100Å. For each complex system, a rigorous protocol of initial energy minimization, thermalization, and a 20 ns conventional MD equilibration phase using the amber forcefield was conducted to ensure system stability. A 9 Å cut-off distance was applied to both van der Waals and short-range electrostatic interactions, while long-range electrostatic interactions were calculated using the particle-mesh Ewald summation method. MD simulations maintained a 2-fs integration time step and employed the SHAKE algorithm for all hydrogen bonds. The lipid tails underwent a focused energy minimization process using the conjugate gradient algorithm, followed by a melting phase at 310 K. Following initial equilibrations, the systems were further stabilized through constant pressure, and temperature (NPT) runs, applying harmonic position restraints to crucial protein and ligand atoms. This preparatory phase culminated in 20 ns conventional MD simulations under consistent conditions, with specific constraints to maintain the bilayer’s integrity.

Gaussian Accelerated Molecular Dynamics (GaMD) provides unconstrained enhanced sampling of molecular configurations and allows retrieval of biomolecular free energies using cumulant expansion[27]. In this work, GaMD simulations were initiated with a 10-ns conventional MD run to gather potential energy statistics and determine GaMD acceleration parameters using AMBER 20. This was followed by a 50-ns equilibration phase upon introducing the boost potential. Eleven independent 800-ns GaMD simulations with randomised initial atomic velocities were conducted for each system.

All GaMD simulations were run at the “dual boost” level by setting the reference energy to the lower bound. Boost potentials were applied to the dihedral energetic term and the total potential energetic term. The upper limit of the boost potential SD, σ0 was set to 6.0 kcal/mol for both the dihedral and the total potential energetic terms.

#### Simulation analysis

All simulations were analysed using VMD [55], distances between Cα atoms of R^3.50^ and E^6.31^ to determine TM3-6 opening and between Cα atoms of Arg95-Arg251-Arg255 to determine triangulation distance for the orthosteric binding pocket. All distances, angles, and RMSF values were determined as averages of simulations.

The *PyReweighting* [56] was used to reweight the distances, the perimeter, and the tilt angle, to compute the PMF profiles. The reweight was calculated using Maclaurin series expansion and a bin size of 0.5Å was used for atom distances, and perimeter, and a bin size of 0.2 Å and 0.2*°* for the tilt angle. The PMF profiles were obtained as averages of simulations in a system.

Graphical visualizations are generated using PyMOL[57] .

### Wet-lab Methods

#### Materials

Dulbeccós Modified Eagle Medium (DMEM) 1885 GlutaMAX™ (GIBCO Cat# 11885-084), penicillin/streptomycin (Substrate Department – UCPH), Trypsin (Bioscience Cat# BE17-161E), phosphate-buffered saline (PBS) (Substrate Department – UCPH), foetal bovine serum (FBS) (Sigma-Aldrich Cat# 12103C), COS 7 cells (ATCC ATTC® CRL-1651), YSi SPA scintillation beards (PerkinElmer Cat# RPNQ0010), Hankś balanced salt solution (HBSS) (GIBCO Cat# 14025050), Sodium succinate dibasic (Sigma-Aldrich Cat# 14160 CAS: 150-90-3).

#### Plasmids

All receptor constructs for murine and human SUCNR1, all mutants, and Gqi4myr were expressed in the pcDNA3.1(+) vector. All receptor constructs have been modified with an N-terminal FLAG-tag and GGGGGGS inserted before the receptor.

#### Cell Culture and Transfection

COS7 was maintained in DMEM 1885 supplemented with 10% FBS 100 units/ml penicillin, and 100 μg/ml streptomycin at 37°C with 10% CO_2_ for COS7 cells.

Cells were transfected using calcium phosphate precipitation with chloroquine addition and supplemented with fresh medium after 5 h.

#### IP accumulation assays

COS7 cells were seeded onto poly-d-lysine-coated 96-well plates at a density of 20,000 cells per well. The following day, cells were transfected with 100 μL transfection medium/well for a total duration of 5 hours. Subsequently, cells were incubated overnight with 0.5 μCi/ml myo [3H] inositol (Perkin Elmer) in a 100 μL growth medium. On the assay day, cells were washed twice with 200 μL HBSS (GIBCO) and were pre-incubated for 5 minutes at 37°C with 100 μL HBSS supplemented with 10 mM LiCl. Succinate addition was followed by a 90-minute incubation at 37°C. Cell lysis was carried out with 40 μL of 10 mM formic acid, followed by a 30-minute incubation on ice. Subsequently, 35 μL of the cell extract was transferred to a white 90-well plate and added 60 μL of 1:8 diluted Ysi SPA scintillation beads (Perkin Elmer). After vigorous shaking for 15 minutes, the plates were centrifuged at 1500 rpm for 5 minutes, and light emission (scintillation) was recorded after a 4-hour delay on a Microbeta (Perkin Elmer). Each determination was performed in triplicates. Gi signaling via IP signaling was mediated through the ‘promiscuous’ G protein Gqi4myr, which binds the receptor-like Gi but mediates Gq signaling[58]

#### BRET and luminescence assay

HEK293 cells were seeded in 6-well plates at a density of 700.000 cells per well. The following day, cells were transfected using lipofectamine 2000 and OptiMEM™ (GIBCO) using varying receptor doses from 1 ng to 300 ng, 500 ng KRAS-venus, and 50 ng of either NES-Nluc-mGsi or -mGsq. An empty vector was co-transfected to balance the change in receptor expression. The following day cells were reseeded into 96-well Nunc white-welled plates at a density of 100.000 cells per well in FluoroBRITE DMEM (GIBCO). On day 4, media was removed and replaced with 80 μL HBSS (GIBCO) containing 5 mM Hepes (GIBCO) and furimazine substrate (Promega). The plates are measured for 8 minutes to obtain a baseline. The BRET signal was calculated from the ratio of the emission intensity at 535 nm to the emission intensity at 475 nm. Each determination was performed in duplicates.

#### Data processing

The findings are presented as the mean ± standard error of the mean (SEM). Using GraphPad Prism software (version 10.0.0 for Windows, GraphPad Software, Boston, Massachusetts USA), we utilized non-linear regression to determine the Top and Bottom values, EC50, and HillSlope of the dose- response curves.

## Reference

1. Hilger, D., M. Masureel, and B.K. Kobilka, Structure and dynamics of GPCR signaling complexes. Nature Structural & Molecular Biology, 2018. 25(1): p. 4–12.

2. Yang, D., et al., G protein-coupled receptors: structure- and function-based drug discovery. Signal Transduction and Targeted Therapy, 2021. 6(1): p. 7.

3. Husted, A.S., et al., GPCR-Mediated Signaling of Metabolites. Cell Metabolism, 2017. 25(4): p. 777–796.

4. Strassheim, D., et al., Metabolite G-Protein Coupled Receptors in Cardio-Metabolic Diseases. Cells, 2021. 10(12): p. 3347.

5. Cosín-Roger, J., et al., Metabolite Sensing GPCRs: Promising Therapeutic Targets for Cancer Treatment? Cells, 2020. 9(11): p. 2345.

6. Recio, C., et al., The Role of Metabolite-Sensing G Protein-Coupled Receptors in Inflammation and Metabolic Disease. Antioxid Redox Signal, 2018. 29(3): p. 237–256.

7. Zhang, X., et al., Structural basis for the ligand recognition and signaling of free fatty acid receptors. Science Advances, 2024. 10(2): p. eadj2384.

8. Li, B., et al., Structural insights into signal transduction of the purinergic receptors P2Y1R and P2Y12R. Protein & Cell, 2022. 14(5): p. 382–386.

9. Li, F., et al., Molecular recognition and activation mechanism of short-chain fatty acid receptors FFAR2/3. Cell Research, 2024. 34(4): p. 323–326.

10. Velcicky, J., et al., Discovery and Optimization of Novel SUCNR1 Inhibitors: Design of Zwitterionic Derivatives with a Salt Bridge for the Improvement of Oral Exposure. Journal of Medicinal Chemistry, 2020. 63(17): p. 9856–9875.

11. Haffke, M., et al., Structural basis of species-selective antagonist binding to the succinate receptor. Nature, 2019. 574(7779): p. 581–585.

12. Shenol, A., et al., Multiple recent HCAR2 structures demonstrate a highly dynamic ligand binding and G protein activation mode. Nature Communications, 2024. 15(1): p. 5364.

13. He, W., et al., Citric acid cycle intermediates as ligands for orphan G-protein-coupled receptors. Nature, 2004. 429(6988): p. 188–93.

14. Reddy, A., et al., pH-Gated Succinate Secretion Regulates Muscle Remodeling in Response to Exercise. Cell, 2020. 183(1): p. 62–75.e17.

15. Löffler, J., et al., A comprehensive molecular profiling approach reveals metabolic alterations that steer bone tissue regeneration. Communications Biology, 2023. 6(1): p. 327.

16. Villanueva-Carmona, T., et al., SUCNR1 signaling in adipocytes controls energy metabolism by modulating circadian clock and leptin expression. Cell Metab, 2023. 35(4): p. 601–619.e10.

17. Keiran, N., et al., SUCNR1 controls an anti-inflammatory program in macrophages to regulate the metabolic response to obesity. Nat Immunol, 2019. 20(5): p. 581–592.

18. Winther, S., M. Trauelsen, and T.W. Schwartz, Protective succinate-SUCNR1 metabolic stress signaling gone bad. Cell Metabolism, 2021. 33(7): p. 1276–1278.

19. Gilissen, J., et al., Insight into SUCNR1 (GPR91) structure and function. Pharmacol Ther, 2016. 159: p. 56–65.

20. Trauelsen, M., et al., Extracellular succinate hyperpolarizes M2 macrophages through SUCNR1/GPR91-mediated Gq signaling. Cell Rep, 2021. 35(11): p. 109246.

21. Rexen Ulven, E., et al., Structure-Activity Investigations and Optimisations of Non-metabolite Agonists for the Succinate Receptor 1. Sci Rep, 2018. 8(1): p. 10010.

22. Trauelsen, M., et al., Receptor structure-based discovery of non-metabolite agonists for the succinate receptor GPR91. Mol Metab, 2017. 6(12): p. 1585–1596.

23. Shenol, A., et al., Molecular dynamics-based identification of binding pathways and two distinct high-affinity sites for succinate in succinate receptor 1/GPR91. Mol Cell, 2024. 84(5): p. 955–966.e4.

24. Hollingsworth, S.A. and R.O. Dror, Molecular Dynamics Simulation for All. Neuron, 2018. 99(6): p. 1129–1143.

25. Wang, T., et al., Molecular activation and G protein coupling selectivity of human succinate receptor SUCR1. Cell Research, 2024.

26. Liu, A., et al., Structural insights into ligand recognition and activation of the succinate receptor SUCNR1. Cell Rep, 2024. 43(7): p. 114381.

27. Miao, Y., V.A. Feher, and J.A. McCammon, Gaussian Accelerated Molecular Dynamics: Unconstrained Enhanced Sampling and Free Energy Calculation. Journal of Chemical Theory and Computation, 2015. 11(8): p. 3584–3595.

28. Heo, L., & Feig, M. Multi-state modeling of G-protein coupled receptors at experimental accuracy. Proteins: Structure, Function, and Bioinformatics, 2022. 90(11), 1873–1885.

29. Jumper, J., et al., Highly accurate protein structure prediction with AlphaFold. Nature, 2021. 596(7873): p. 583–589.

30. Li, Y., et al., The full activation mechanism of the adenosine A(1) receptor revealed by GaMD and Su-GaMD simulations. Proc Natl Acad Sci U S A, 2022. 119(42): p. e2203702119.

31. Do, H.N., J. Wang, and Y. Miao, Deep Learning Dynamic Allostery of G-Protein-Coupled Receptors. JACS Au, 2023. 3(11): p. 3165–3180.

32. Madhu, M.K., K. Shewani, and R.K. Murarka, Biased Signaling in Mutated Variants of β2- Adrenergic Receptor: Insights from Molecular Dynamics Simulations. Journal of Chemical Information and Modeling, 2024. 64(2): p. 449–469.

33. Conrad, M., et al., Agonist Binding and G Protein Coupling in Histamine H(2) Receptor: A Molecular Dynamics Study. Int J Mol Sci, 2020. 21(18).

34. Wang, J., et al., Gaussian accelerated molecular dynamics (GaMD): principles and applications. Wiley Interdiscip Rev Comput Mol Sci, 2021. 11(5).

35. Sandhu, M., et al., Conformational plasticity of the intracellular cavity of GPCR−G-protein complexes leads to G-protein promiscuity and selectivity. Proceedings of the National Academy of Sciences, 2019. 116(24): p. 11956–11965.

36. Wheatley, M., et al., Lifting the lid on GPCRs: the role of extracellular loops. Br J Pharmacol, 2012. 165(6): p. 1688–1703.

37. Hauser, A.S., et al., GPCR activation mechanisms across classes and macro/microscales. Nature Structural & Molecular Biology, 2021. 28(11): p. 879–888.

38. Xu, X., et al., Binding pathway determines norepinephrine selectivity for the human β(1)AR over β(2)AR. Cell Res, 2021. 31(5): p. 569–579.

39. Miao, Y. and J.A. McCammon, Graded activation and free energy landscapes of a muscarinic G- protein-coupled receptor. Proc Natl Acad Sci U S A, 2016. 113(43): p. 12162–12167.

40. Els, S., A.G. Beck-Sickinger, and C. Chollet, Ghrelin receptor: high constitutive activity and methods for developing inverse agonists. Methods Enzymol, 2010. 485: p. 103–21.

41. Canals, M. and G. Milligan, Constitutive Activity of the Cannabinoid CB1 Receptor Regulates the Function of Co-expressed Mu Opioid Receptors. Journal of Biological Chemistry, 2008. 283(17): p. 11424–11434.

42. Latorraca, N.R., A.J. Venkatakrishnan, and R.O. Dror, GPCR Dynamics: Structures in Motion. Chemical Reviews, 2017. 117(1): p. 139–155.

43. Miao, Y., et al., Activation and dynamic network of the M2 muscarinic receptor. Proc Natl Acad Sci U S A, 2013. 110(27): p. 10982–7.

44. Miao, Y., A.D. Caliman, and J.A. McCammon, Allosteric effects of sodium ion binding on activation of the m3 muscarinic g-protein-coupled receptor. Biophys J, 2015. 108(7): p. 1796–1806.

45. Dror, R.O., et al., Activation mechanism of the β2-adrenergic receptor. Proc Natl Acad Sci U S A, 2011. 108(46): p. 18684–9.

46. Xu, X., et al., Constrained catecholamines gain β2AR selectivity through allosteric effects on pocket dynamics. Nature Communications, 2023. 14(1): p. 2138.

47. Manglik, A., et al., *Structural Insights into the Dynamic Process of &#x3b*2*; 2- Adrenergic Receptor Signaling*. Cell, 2015. 161(5): p. 1101–1111.

48. Wu, Y., et al., MD Simulations Revealing Special Activation Mechanism of Cannabinoid Receptor 1. Frontiers in Molecular Biosciences, 2022. 9.

49. Abramson, J., et al., Accurate structure prediction of biomolecular interactions with AlphaFold 3. Nature, 2024.

50. 2023-4, S.R., Protein Preparation Wizard; Epik, Schrödinger, LLC, New York, NY,2023; Impact, Schrödinger, LLC, New York, NY; Prime, Schrödinger, LLC, New York, NY, 2023. 2023-4.

51. Schrödinger Release 2023-4: Glide, S., LLC, New York, NY, 2024., 2023.

52. Jo, S., et al., CHARMM-GUI: A web-based graphical user interface for CHARMM. Journal of Computational Chemistry, 2008. 29(11): p. 1859–1865.

53. Lomize, M.A., et al., OPM database and PPM web server: resources for positioning of proteins in membranes. Nucleic Acids Research, 2011. 40(D1): p. D370–D376.

54. Tian, C., et al., ff19SB: Amino-Acid-Specific Protein Backbone Parameters Trained against Quantum Mechanics Energy Surfaces in Solution. Journal of Chemical Theory and Computation, 2020. 16(1): p. 528–552.

55. Humphrey, W., A. Dalke, and K. Schulten, VMD: visual molecular dynamics. J Mol Graph, 1996. 14(1): p. 33–8, 27-8.

56. Miao, Y., et al., Improved Reweighting of Accelerated Molecular Dynamics Simulations for Free Energy Calculation. Journal of Chemical Theory and Computation, 2014. 10(7): p. 2677–2689.

57. The PyMOL Molecular Graphics System, Version 3.0 Schrödinger, LLC. Kostenis, E., F.Y. Zeng, and J. Wess, Functional characterization of a series of mutant G protein alphaq subunits displaying promiscuous receptor coupling properties. J Biol Chem, 1998. 273(28): p. 17886–92.

