## Supplementary figures for "Constitutive activation and allosteric mechanisms underlying Gi/Gq signaling bias in SUCNR1 revealed by AlphaFold-based modeling and enhanced sampling simulations"

Table1

| # | Specie | Structure | Length of simulation | TM3-6 opening | DRY motif – $R^{3.50}$ shift upwards | NPxxY motif – $Tyr^{7.58}$ translation towards DRY | ECL2 - opening | R248 – final position |
|---|---|---|---|---|---|---|---|---|
| 1 | Rat | PDB:6ibb | 800ns | No | No | No | No | Between Y171 and Y103 |
| 2 | Rat | PDB:6ibb | 800ns | No | No | No | No | Between Y171 and Y103 |
| 3 | Rat | PDB:6ibb | 800ns | No | No | No | No | Between Y171 and Y103 |
| 4 | Rat | PDB:6ibb | 800ns | No | No | No | No | Between Y171 and Y103 |
| 5 | Rat | PDB:6ibb | 800ns | No | No | No | No | Between Y171 and Y103 |
| 6 | Mouse | AlphaFold | 800ns | Yes | No | No | No | Between Y171 and Y103 |
| 7 | Mouse | AlphaFold | 800ns | Yes | No | No | No | Between Y171 and Y103 |
| 8 | Mouse | AlphaFold | 800ns | No | No | No | Yes | Between Y171 and Y103 |
| 9 | Mouse | AlphaFold | 800ns | No | No | No | No | Between Y171 and Y103 |
| 10 | Mouse | AlphaFold | 800ns | No | No | No | Yes | Shortly interacts with D174 |
| 11 | Mouse | AlphaFold | 800ns | No | No | No | No | Between Y171 and Y103 |
| 12 | Mouse | AlphaFold | 800ns | No | No | No | No | Between Y171 and Y103 |
| 13 | Mouse | AlphaFold | 800ns | No | No | No | No | Between Y171 and Y103 |
| 14 | Mouse | AlphaFold | 800ns | No | No | No | Yes | Shortly interacts with D174 |
| 15 | Mouse | AlphaFold | 800ns | No | No | No | No | Between Y171 and Y103 |
| 16 | Mouse | AlphaFold | 800ns | No | No | No | Yes | Between Y171 and Y103 |
| 17 | Human | AlphaFold | 800ns | Yes | Yes | No | Yes | Between F175 and Y107 |
| 18 | Human | AlphaFold | 800ns | Yes | Yes | No | Yes | Interacts with D174 |
| 19 | Human | AlphaFold | 800ns | No | Yes | No | Yes | Between F175 and Y107 |
| 20 | Human | AlphaFold | 800ns | Yes | Yes | No | Yes | Between F175 and Y107 |
| 21 | Human | AlphaFold | 800ns | Yes | Yes | Yes | Yes | Between F175 and Y107 |
| 22 | Human | AlphaFold | 800ns | No | Yes | Yes | Yes | Between F175 and Y107 |
| 23 | Human | AlphaFold | 800ns | No | Yes | Yes | Yes | Between F175 and Y107 |
| 24 | Human | AlphaFold | 800ns | Yes | Yes | No | Yes | Interacts with D174 |
| 25 | Human | AlphaFold | 800ns | Yes | Yes | No | Yes | Between F175 and Y107 |
| 26 | Human | AlphaFold | 800ns | Yes | Yes | Yes | Yes | Between F175 and Y107 |
| 26 | Human | AlphaFold | 800ns | yes | Yes | No | Yes | Between F175 and Y107 |

**Table 1. All simulations of SUCNR1 without a G-protein and without a ligand, for three different species (rat, mouse and human).** TM3-6 is considered open for distance above 14 Å . Next, we noted the position of $R^{3.50}$ in upwards conformation and the transition of $Y^{7.58}$ towards it (landmarks of GPCR activation) . ECL2 was classified as open for distances between $D^{45.52}$ and $Y^{7.35}$ above 7Å . The final position of R248 in respect to its interaction partners is noted in the last column.

Table 2

| # | Specie | Structure | Length of simulation | Receptor coupling | TM3-6 opening | DRY motif – $R^{3.50}$ shift upwards | NPxxY motif – $Tyr^{7.58}$ translation towards DRY | ECL2 - opening | R248 – final position |
|---|---|---|---|---|---|---|---|---|---|
| 1 | Mouse | AlphaFold | 800ns | Gi | Yes | Yes | Yes | Yes | Between Y171 and Y103 |
| 2 | Mouse | AlphaFold | 800ns | Gi | Yes | Yes | Yes | Yes | Between Y171 and Y103 |
| 3 | Mouse | AlphaFold | 800ns | Gi | Yes | Yes | Yes | Yes | Shortly interacts with D170 |
| 4 | Mouse | AlphaFold | 800ns | Gi | Yes | Yes | Yes | Yes | Between Y171 and Y103 |
| 5 | Mouse | AlphaFold | 800ns | Gi | Yes | Yes | Yes | Yes | Between Y171 and Y103 |
| 6 | Mouse | AlphaFold | 800ns | Gi | Yes | Yes | Yes | Yes | Between Y171 and Y103 |
| 7 | Mouse | AlphaFold | 800ns | Gi | Yes | Yes | Yes | Yes | Between Y171 and Y103 |
| 8 | Mouse | AlphaFold | 800ns | Gi | Yes | Yes | Yes | Yes | Between Y171 and Y103 |
| 9 | Mouse | AlphaFold | 800ns | Gi | Yes | Yes | Yes | Yes | Between Y171 and Y103 |
| 10 | Mouse | AlphaFold | 800ns | Gi | Yes | Yes | Yes | Yes | Between Y171 and Y103 |
| 11 | Mouse | AlphaFold | 800ns | Gi | Yes | Yes | Yes | Yes | Shortly interacts with D170 |
| 12 | Mouse | AlphaFold | 800ns | Gq | Yes | Yes | Yes | Yes | Between Y171 and Y103 |
| 13 | Mouse | AlphaFold | 800ns | Gq | Yes | Yes | Yes | Yes | Between Y171 and Y103 |
| 14 | Mouse | AlphaFold | 800ns | Gq | Yes | Yes | Yes | Yes | Shortly interacts with D170 |
| 15 | Mouse | AlphaFold | 800ns | Gq | Yes | Yes | Yes | Yes | Between Y171 and Y103 |
| 16 | Mouse | AlphaFold | 800ns | Gq | Yes | Yes | Yes | Yes | Shortly interacts with D170 |
| 17 | Mouse | AlphaFold | 800ns | Gq | Yes | Yes | Yes | Yes | Between Y171 and Y103 |
| 18 | Mouse | AlphaFold | 800ns | Gq | Yes | Yes | Yes | Yes | Between Y171 and Y103 |
| 19 | Mouse | AlphaFold | 800ns | Gq | Yes | Yes | Yes | Yes | Interacts wth D170 |
| 20 | Mouse | AlphaFold | 800ns | Gq | Yes | Yes | Yes | Yes | Shortly interacts with D170 |
| 21 | Mouse | AlphaFold | 800ns | Gq | Yes | Yes | Yes | Yes | Between Y171 and Y103 |
| 22 | Mouse | AlphaFold | 800ns | Gq | Yes | Yes | Yes | Yes | Between Y171 and Y103 |
| 23 | Human | AlphaFold | 800ns | Gi | Yes | Yes | Yes | Yes | Interacts with D174 |
| 24 | Human | AlphaFold | 800ns | Gi | Yes | Yes | Yes | Yes | Interacts with D174 |
| 25 | Human | AlphaFold | 800ns | Gi | Yes | Yes | Yes | Yes | Interacts with D174 |
| 26 | Human | AlphaFold | 800ns | Gi | Yes | Yes | Yes | Yes | Between F175 and Y107 |
| 27 | Human | AlphaFold | 800ns | Gi | Yes | Yes | Yes | Yes | Between F175 and Y107 |
| 28 | Human | AlphaFold | 800ns | Gi | Yes | Yes | Yes | Yes | Interacts with D174 |
| 29 | Human | AlphaFold | 800ns | Gi | Yes | Yes | Yes | Yes | Interacts with D174 |
| 30 | Human | AlphaFold | 800ns | Gi | Yes | Yes | Yes | Yes | Interacts with D174 |
| 31 | Human | AlphaFold | 800ns | Gi | Yes | Yes | Yes | Yes | Between F175 and Y107 |
| 32 | Human | AlphaFold | 800ns | Gi | Yes | Yes | Yes | Yes | Between F175 and Y107 |
| 33 | Human | AlphaFold | 800ns | Gi | Yes | Yes | Yes | Yes | Interacts with D174 |
| 34 | Human | AlphaFold | 800ns | Gq | Yes | Yes | Yes | Yes | Interacts with D174 |
| 35 | Human | AlphaFold | 800ns | Gq | Yes | Yes | Yes | Yes | Interacts with D174 |
| 36 | Human | AlphaFold | 800ns | Gq | Yes | Yes | Yes | Yes | Interacts with D174 |
| 37 | Human | AlphaFold | 800ns | Gq | Yes | Yes | Yes | Yes | Interacts with D174 |
| 38 | Human | AlphaFold | 800ns | Gq | Yes | Yes | Yes | Yes | Interacts with D174 |
| 39 | Human | AlphaFold | 800ns | Gq | Yes | Yes | Yes | Yes | Interacts with D174 |
| 40 | Human | AlphaFold | 800ns | Gq | Yes | Yes | Yes | Yes | Between F175 and Y107 |
| 41 | Human | AlphaFold | 800ns | Gq | Yes | Yes | Yes | Yes | Interacts with D174 |
| 42 | Human | AlphaFold | 800ns | Gq | Yes | Yes | Yes | Yes | Interacts with D174 |
| 43 | Human | AlphaFold | 800ns | Gq | Yes | Yes | Yes | Yes | Interacts with D174 |
| 44 | Human | AlphaFold | 800ns | Gq | Yes | Yes | Yes | Yes | Between F175 and Y107 |

**Table 2. All simulations of SUCNR1 without a Gi or Gq-protein and without a ligand, in four different systems (hSUCNR1-Gi, hSUCNR1-Gq, mSICNR1-Gi and mSUCNR1-Gq).** TM3-6 is considered open for distance above 14 Å . The position of $R^{3.50}$ in upwards conformation and the transition of $Y^{7.58}$ towards it are noted (landmarks of GPCR activation) . ECL2 was classified as open for distances between $D^{45.52}$ and $Y^{7.35}$ above 7Å. The final position of R248 in respect to its interaction partners is noted in the last column.

### Table 3

| # | Specie | Structure | Length of simulation | Receptor coupling | TM3-6 opening | DRY motif – $R^{3.50}$ shift upwards | NPxxY motif – $Tyr^{7.58}$ translation towards DRY | ECL2 - opening | R248 – final position |
|---|---|---|---|---|---|---|---|---|---|
| 1 | Mouse | AlphaFold | 800ns | Gi | Yes | Yes | Yes | Yes | Intearcts with succinate |
| 2 | Mouse | AlphaFold | 800ns | Gi | Yes | Yes | Yes | Yes | Intearcts with succinate |
| 3 | Mouse | AlphaFold | 800ns | Gi | Yes | Yes | Yes | Yes | Intearcts with succinate |
| 4 | Mouse | AlphaFold | 800ns | Gi | Yes | Yes | Yes | Yes | Intearcts with succinate |
| 5 | Mouse | AlphaFold | 800ns | Gi | Yes | Yes | Yes | Yes | Intearcts with succinate |
| 6 | Mouse | AlphaFold | 800ns | Gq | Yes | Yes | Yes | Yes | Intearcts with succinate |
| 7 | Mouse | AlphaFold | 800ns | Gq | Yes | Yes | Yes | Yes | Intearcts with succinate |
| 8 | Mouse | AlphaFold | 800ns | Gq | Yes | Yes | Yes | Yes | Intearcts with succinate |
| 9 | Mouse | AlphaFold | 800ns | Gq | Yes | Yes | Yes | Yes | Intearcts with succinate |
| 10 | Mouse | AlphaFold | 800ns | Gq | Yes | Yes | Yes | Yes | Intearcts with succinate |
| 11 | Human | AlphaFold | 800ns | Gi | Yes | Yes | Yes | Yes | Intearcts with succinate |
| 12 | Human | AlphaFold | 800ns | Gi | Yes | Yes | Yes | Yes | Intearcts with succinate |
| 13 | Human | AlphaFold | 800ns | Gi | Yes | Yes | Yes | Yes | Intearcts with succinate |
| 14 | Human | AlphaFold | 800ns | Gi | Yes | Yes | Yes | Yes | Intearcts with succinate |
| 15 | Human | AlphaFold | 800ns | Gi | Yes | Yes | Yes | Yes | Intearcts with succinate |
| 16 | Human | AlphaFold | 800ns | Gq | Yes | Yes | Yes | Yes | Intearcts with succinate |
| 17 | Human | AlphaFold | 800ns | Gq | Yes | Yes | Yes | Yes | Intearcts with succinate |
| 18 | Human | AlphaFold | 800ns | Gq | Yes | Yes | Yes | Yes | Intearcts with succinate |
| 19 | Human | AlphaFold | 800ns | Gq | Yes | Yes | Yes | Yes | Intearcts with succinate |
| 20 | Human | AlphaFold | 800ns | Gq | Yes | Yes | Yes | Yes | Intearcts with succinate |

**Table 3. All simulations of SUCNR1 without a Gi or Gq-protein and with succinate in the orthosteric pocket, in four different systems (hSUCNR1-Gi, hSUCNR1-Gq, mSICNR1-Gi and mSUCNR1-Gq).** TM3-6 is considered open for distance above 14 Å . The position of $R^{3.50}$ in upwards conformation and the transition of $Y^{7.58}$ towards it are noted (landmarks of GPCR activation) . ECL2 was classified as open for distances between $D^{45.52}$ and $Y^{7.35}$ above 7Å . The final position of R248 in respect to its interaction partners is noted in the last column.

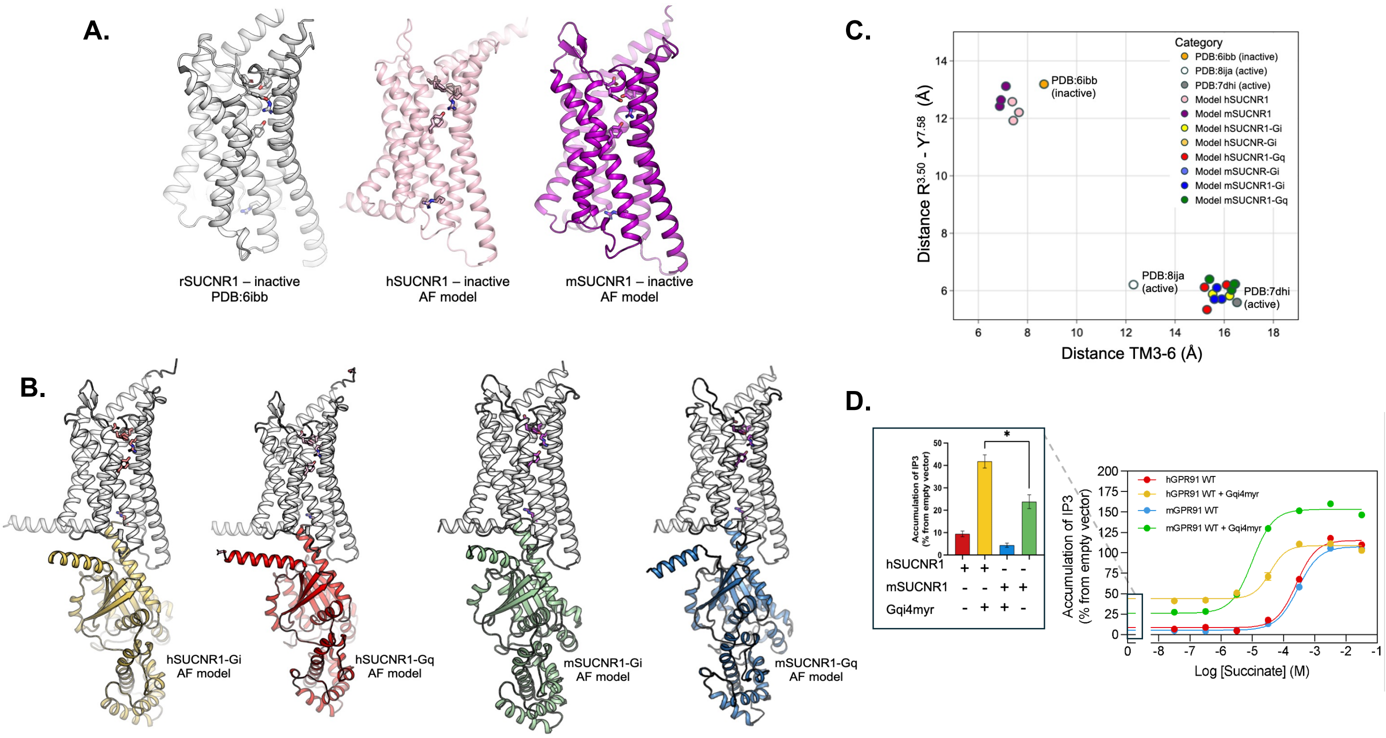


**Supplementary figure 1. Structures used for GAMD simulations**

**A.** Structures used in Apo systems – rSUCNR1 (grey – pdb:6rnk), hSUCNR1 (pink), mSUCNR1(purple). **B.** Structures used in G-coupled systems – hSUCNR1-Gi (yellow), hSUCNR1-Gq (red), mSUCNR1-Gi (green), mSUCNR1-Gq (blue).

**C**. Alpha Fold prediction od all generated structures in realtion to refernces structures. **D** Dose-response curves of mSUCNR and hSUCNR1 when stimulated with Succinate in Gi and Gq signaling pathways. Panel to the left shows the basal activity levels

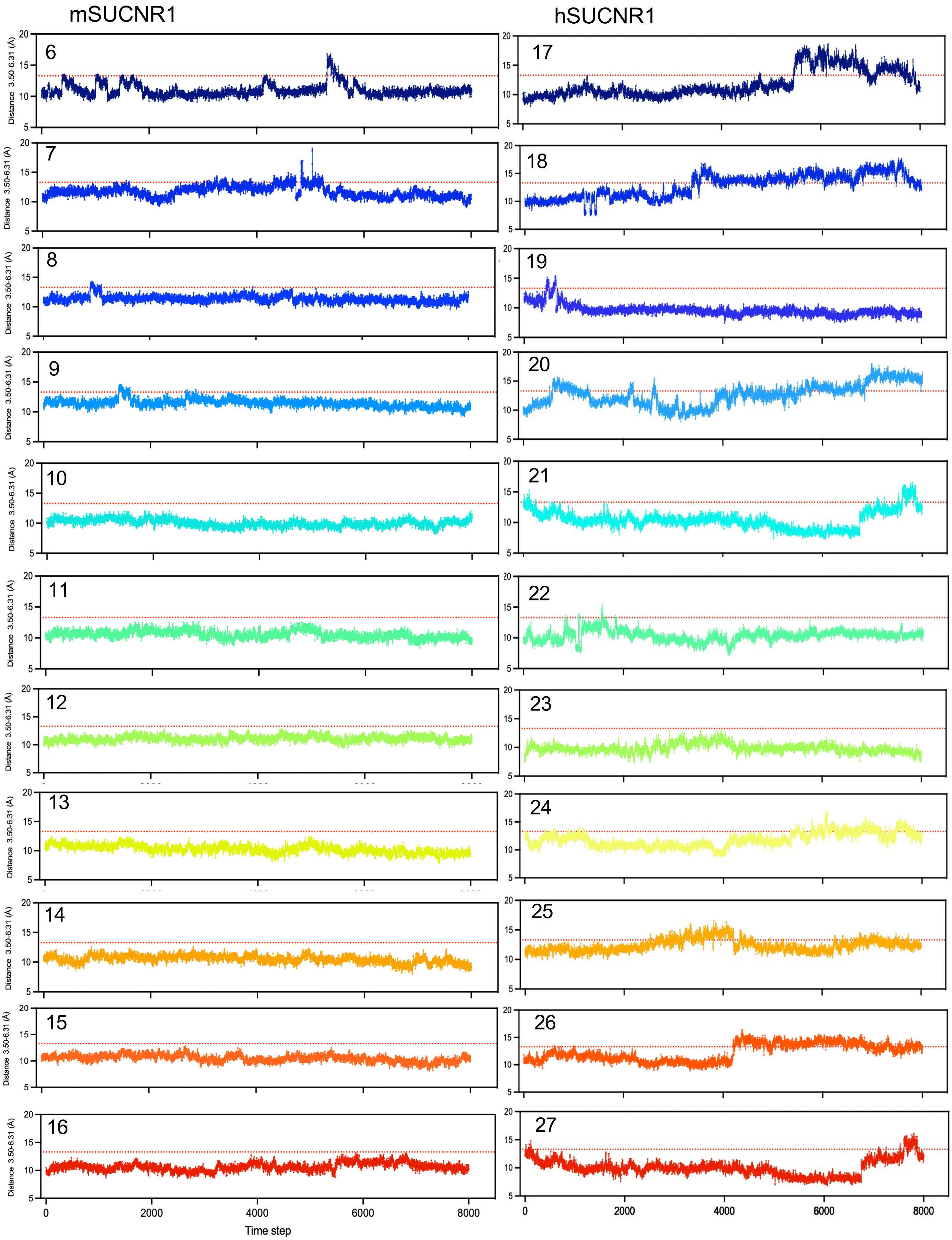


**Supplementary Figure 2.** Distance between $R^{3.50}$ (DRY) – $E^{6.31}$ as a measure for the opening of the TMIII – TMVI in mSUCNR1 and hSUCNR1

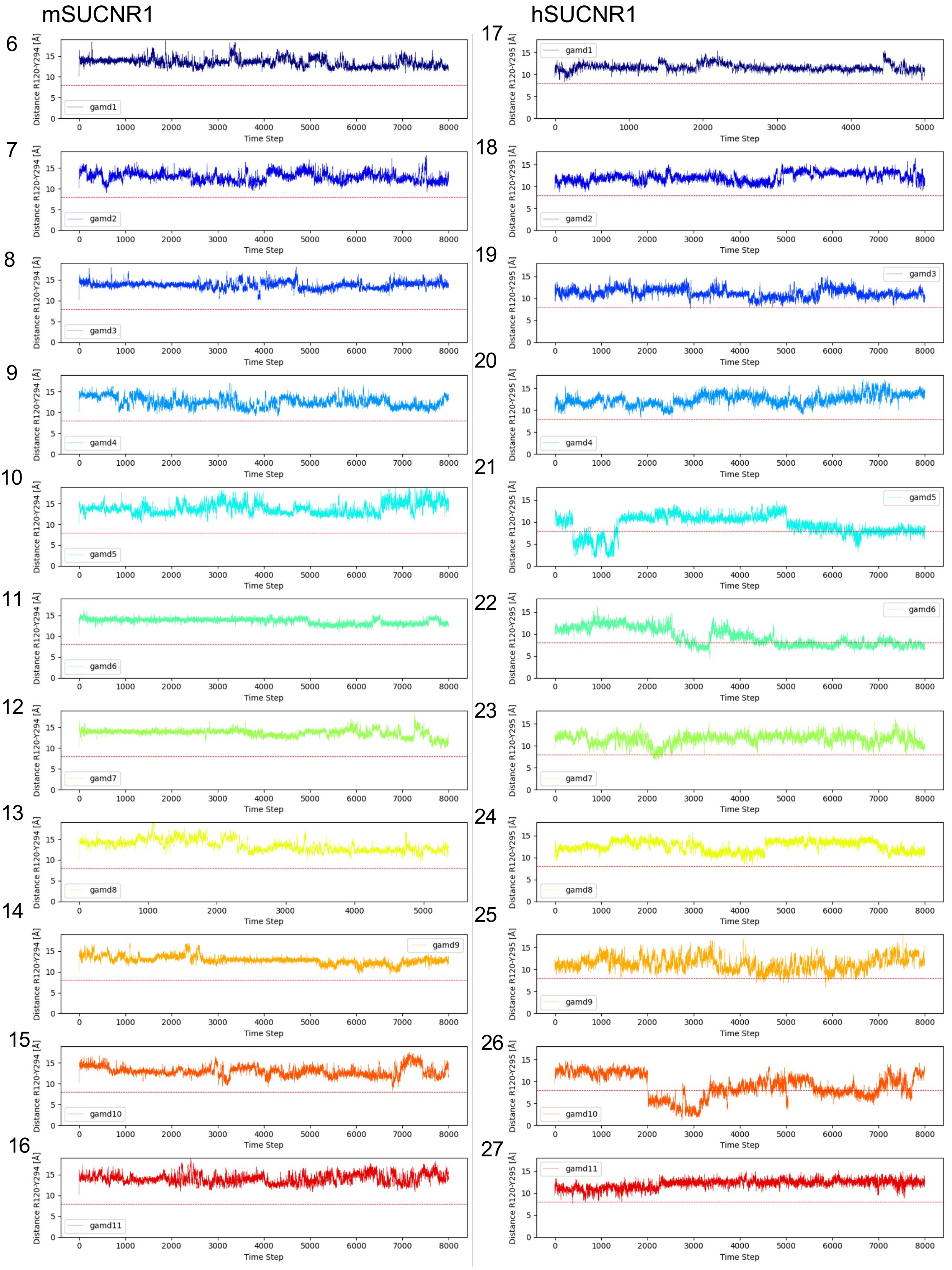


**Supplementary Figure 3.** Distance between $R^{3.50}$ (DRY) – $Y^{7.53}$ (NPxxY) as a measure for the rearrangement of DRY and NPxxY motif for all simulations in apo- mSUCNR1 and hSUCNR1

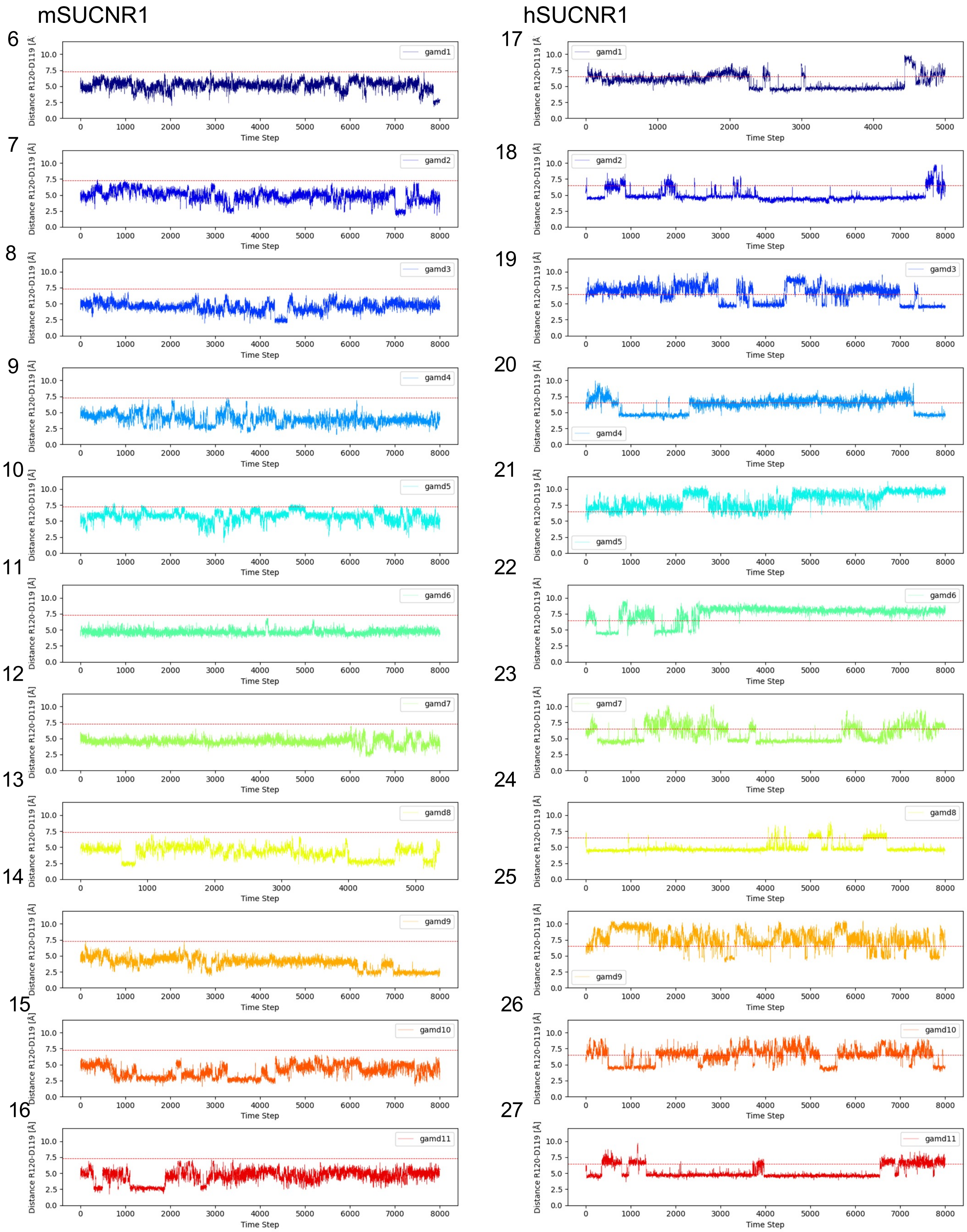


**Supplementary Figure 4.** Distance between $R^{3.50}$ (DRY) – $D^{3.49}$ (DRY) as a measure for the rearrangement of DRY and NPxxY motif for all simulations in apo- mSUCNR1 and hSUCNR1

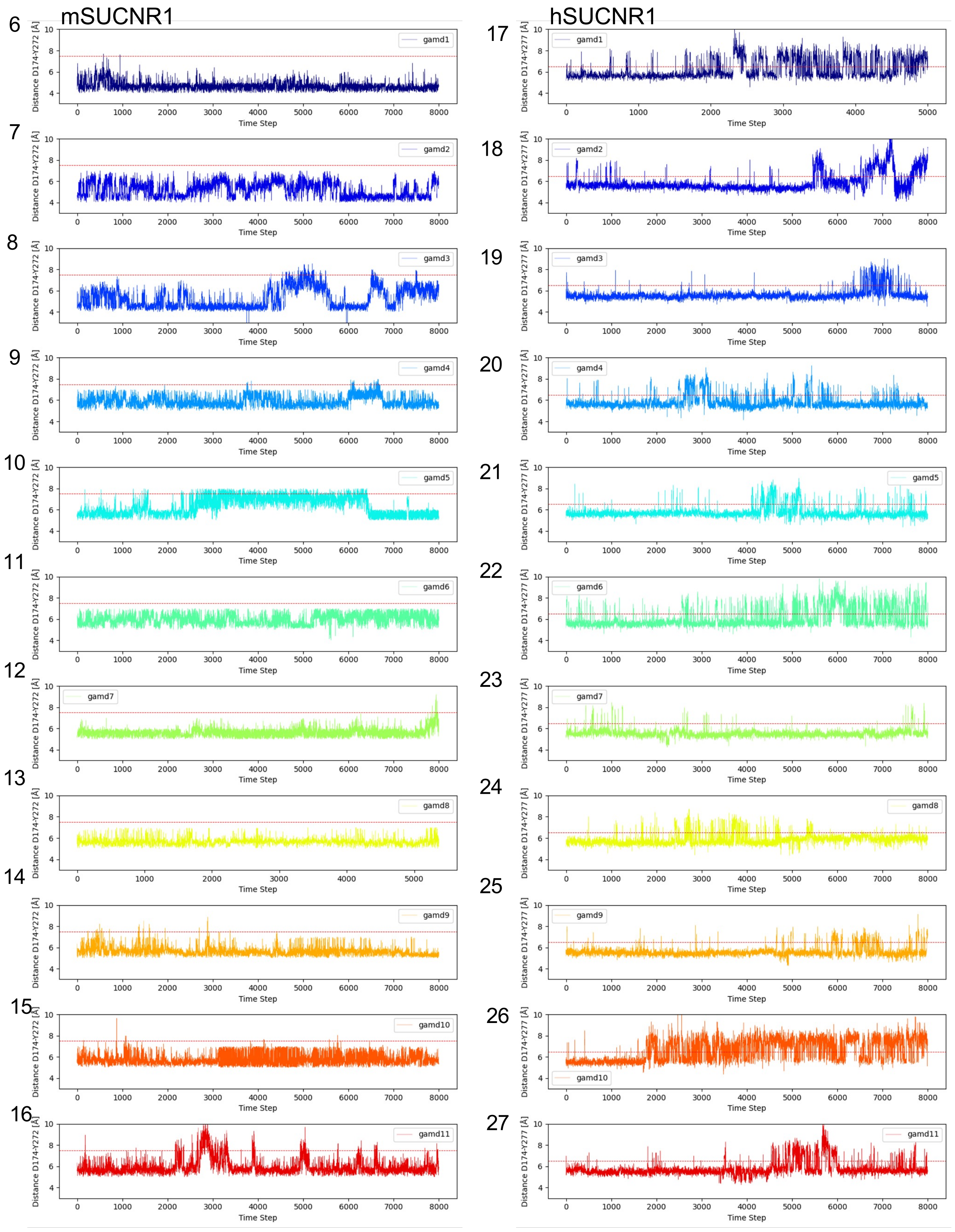


**Supplementary Figure 5.** Distance between $Y^{7.35} – D^{45.52}$ as a measure for the opening of ECl2 for all simulations in apo-mSUCNR1 and hSUCNR1

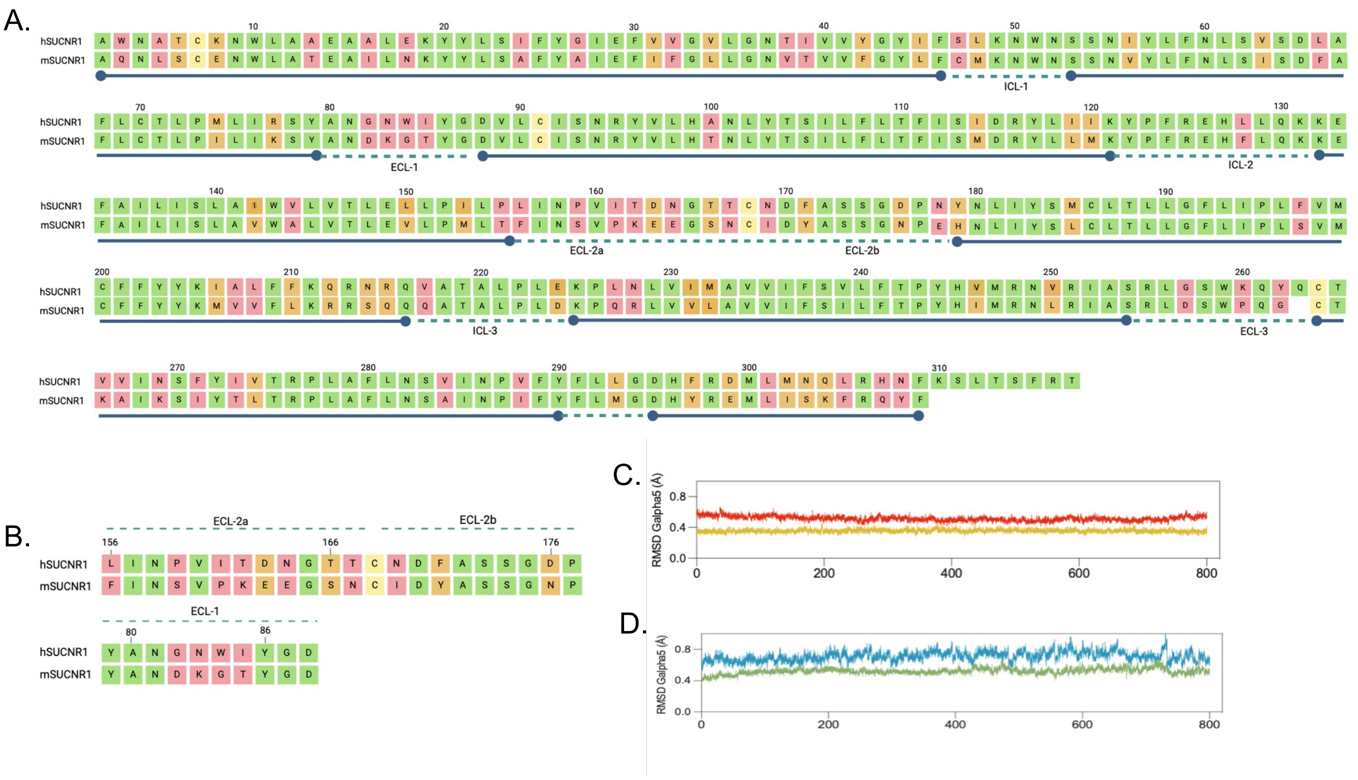


**Supplementary Figure 6. A**. Sequence alignment between hSUCNR1 and mSUCNR1 with similar residues indicated in green, similar residues in yellow and different residues in pink. **B.** Significant differences in the sequence between human in murine SUCNR1 in ECL1 and ECL2 highlighter.**(C-D)** Root mean square deviation (RMSD) of the Gα5 helix when bound to hSUCNR1 (C) and mSUCNR1 (D). Higher RMSD values are observed for Gq compared to Gi, highlighting greater Gα5 helix flexibility for Gq proteins. Average values are shown as bar graphs on the right.

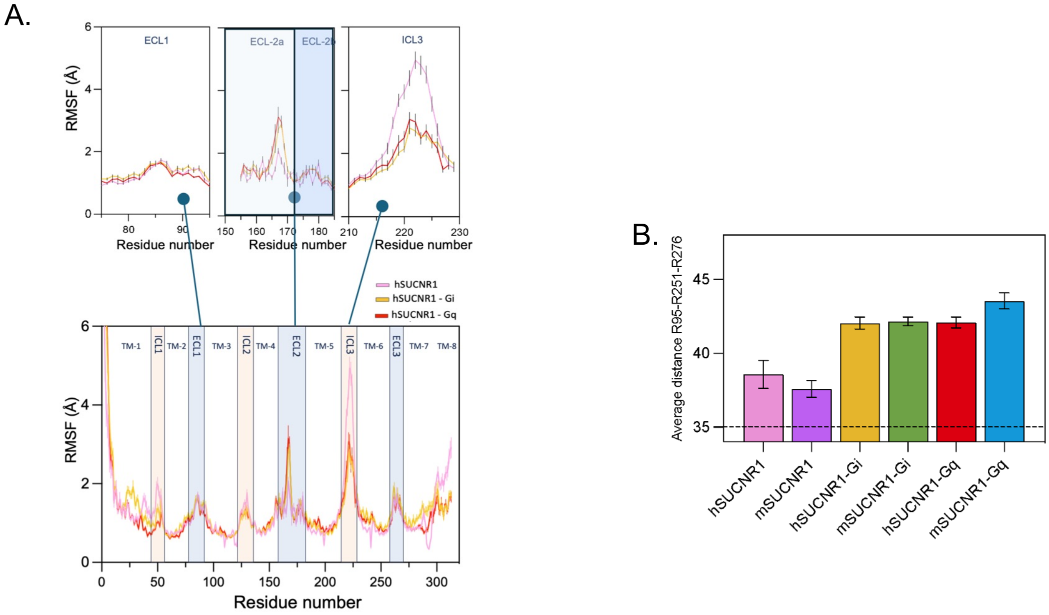


**Supplementary Figure 7. RMSF changes of GPCRs. A**. RMSF of apo-hSUCNR1 (pink), hSUCNR1-Gi (yellow), hSUCNR1-Gq (red), showing an increase in the ECL2 RMSF upon coupling to a G-protein, compared to apo hSUCNR1 and decrease in the ICL3 RMSF for the G-protein coupled, compared to apo hSUCNR1. **B.** Bar plot of the average triangulation distances between arginines. Error bars represent SD.

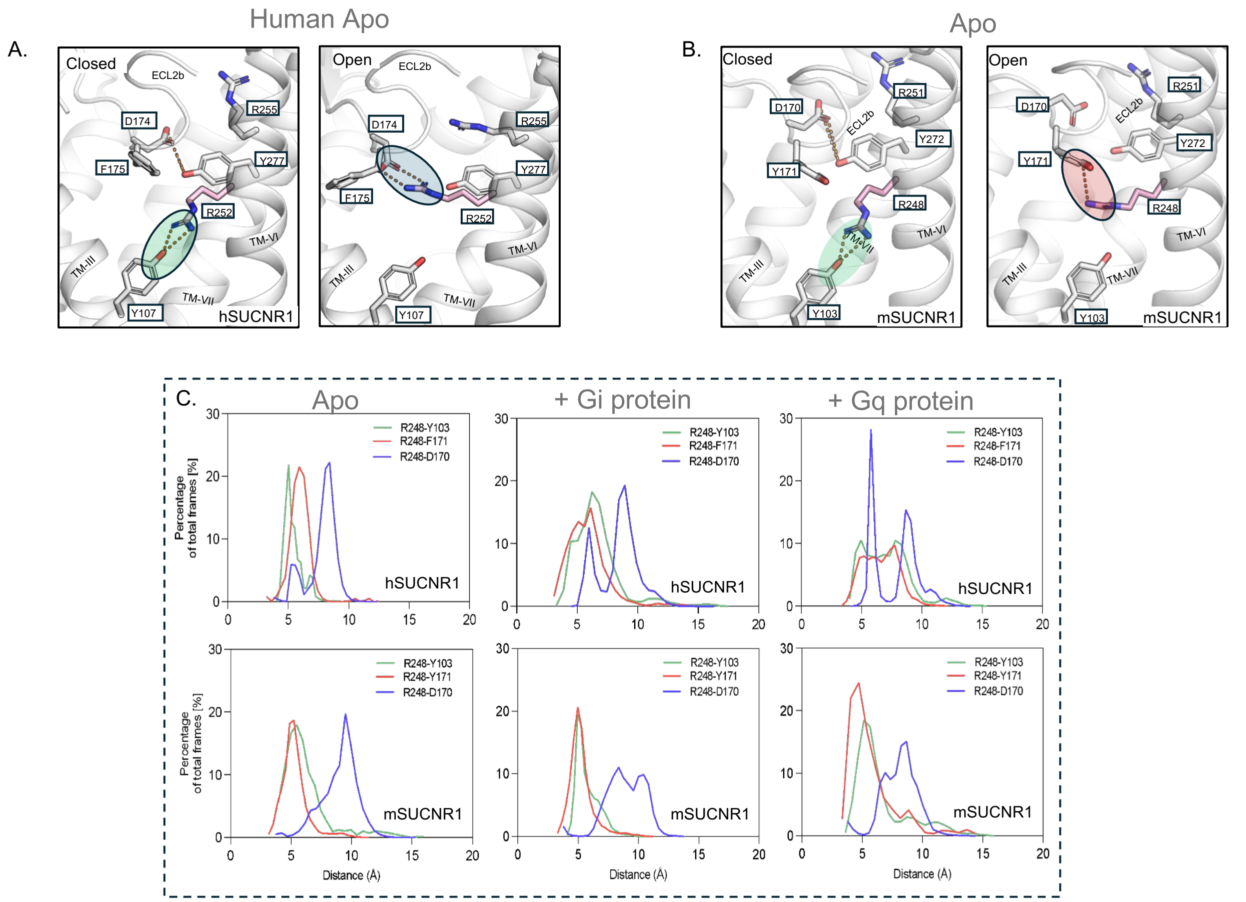


**Supplementary Figure 8: Role of R248 (R6.55) in SUCNR1 Activation**

**A:** In hSUCNR1, the inactive state shows ECL-2 closed, with $R^{6.55}$ interacting with $Y^{3.37}$ and $F^{45.53}$. In the active state, ECL-2 opens, and $R^{6.55}$ flips upward toward the orthosteric pocket, bypassing $F^{45.53}$ to interact with $D^{45.52}$.

**B:** In mSUCNR1, the inactive state features ECL-2 closed, with $R^{6.55}$ interacting with $Y^{3.37}$ and $Y^{45.53}$. In the active state, despite ECL-2 opening, $R^{6.55}$ remains trapped between $Y^{3.37}$ and $Y^{45.53}$, unable to interact with $D^{45.52}$.

**C:** Distance distributions for $R^{6.55}$ interactions in hSUCNR1 and mSUCNR1 reveal species-specific dynamics. In hSUCNR1, a second peak in the G-coupled system confirms $R^{6.55}$ forming a bond with $D^{45.52}$, while distances to Y3.37 and $F^{45.53}$ increase. In mSUCNR1, the $R^{6.55}$-$D^{45.52}$ distance shortens slightly in the G-coupled state but fails to establish a distinct peak, with interactions with $Y^{3.37}$ and $Y^{45.53}$ remaining stable.

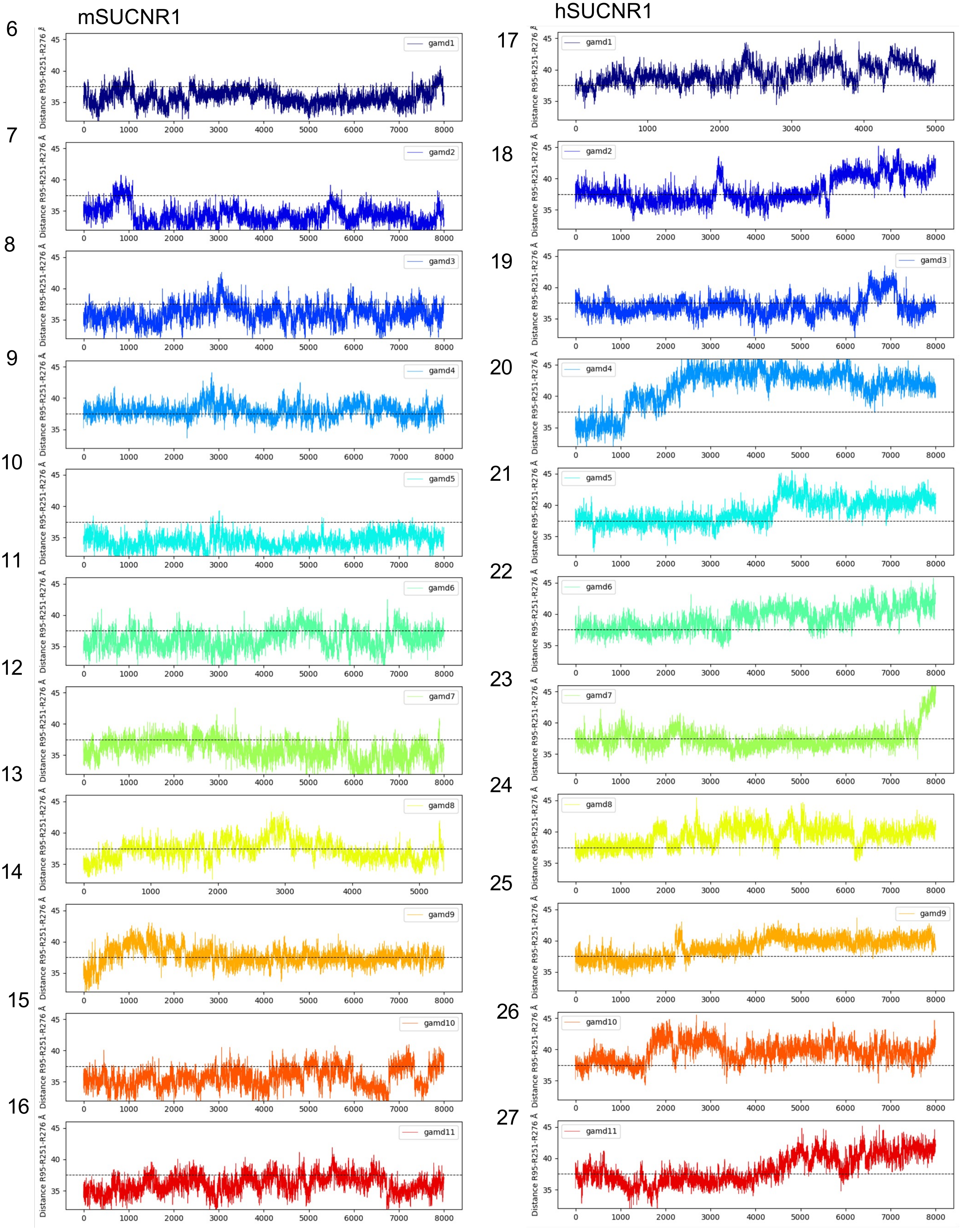


**Supplementary Figure 9.** Triangulation distance between R$^{3.29}$-R$^{6.58}$-R$^{7.39}$ as a measure for the size of the orthosteric site in the apo-systems of mSUCNR1 and hSUCNR1 for all simulations

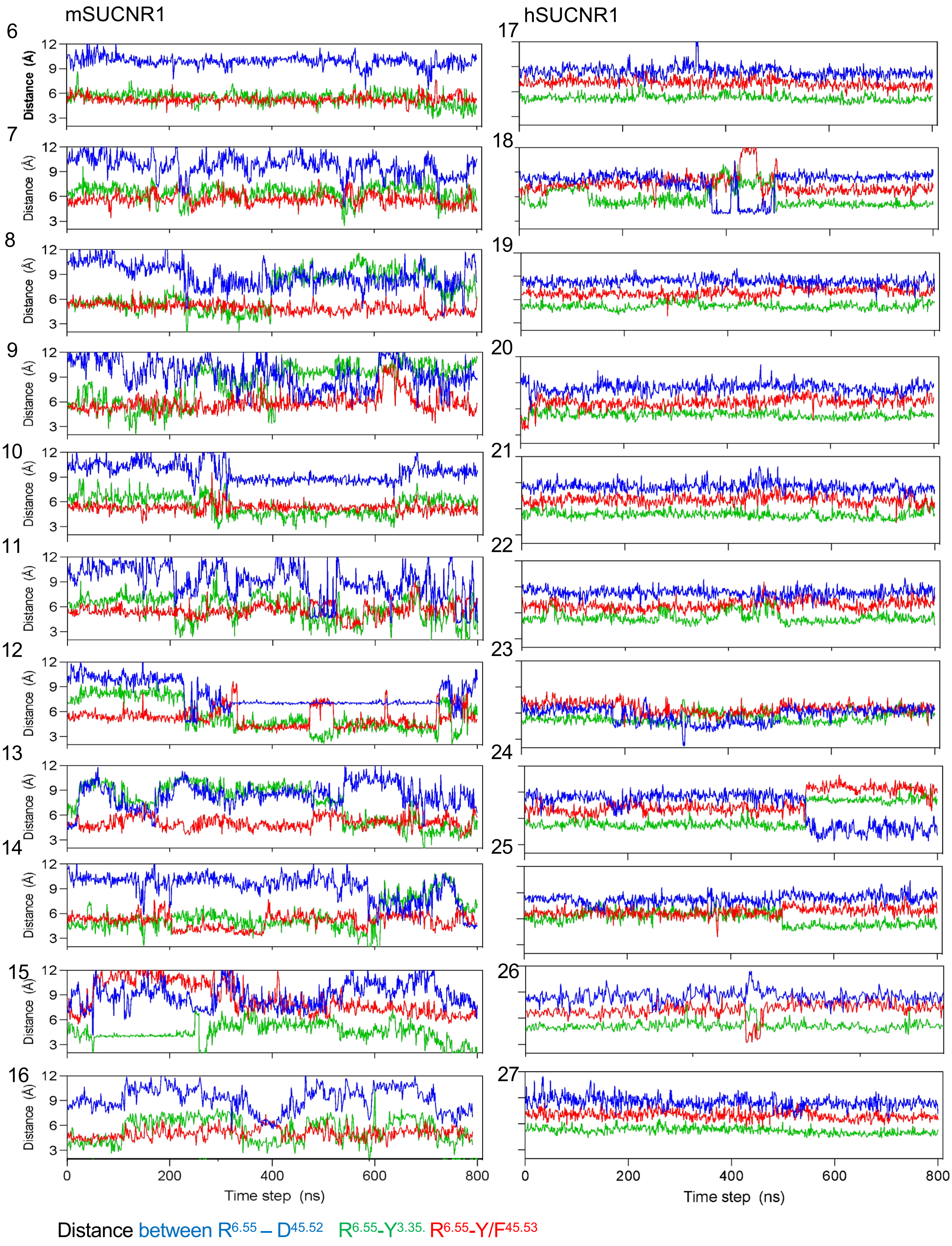


**Supplementary Figure 10.** Distance between $R^{6.55}$ – $D^{45.52}$ (blue), $R^{6.55}$-$Y^{3.35}$ (green), $R^{6.55}$-$Y/F^{45.53}$ (red) through every simulation **A**. – mSUCNR1 and **B.** - hSUCNR1

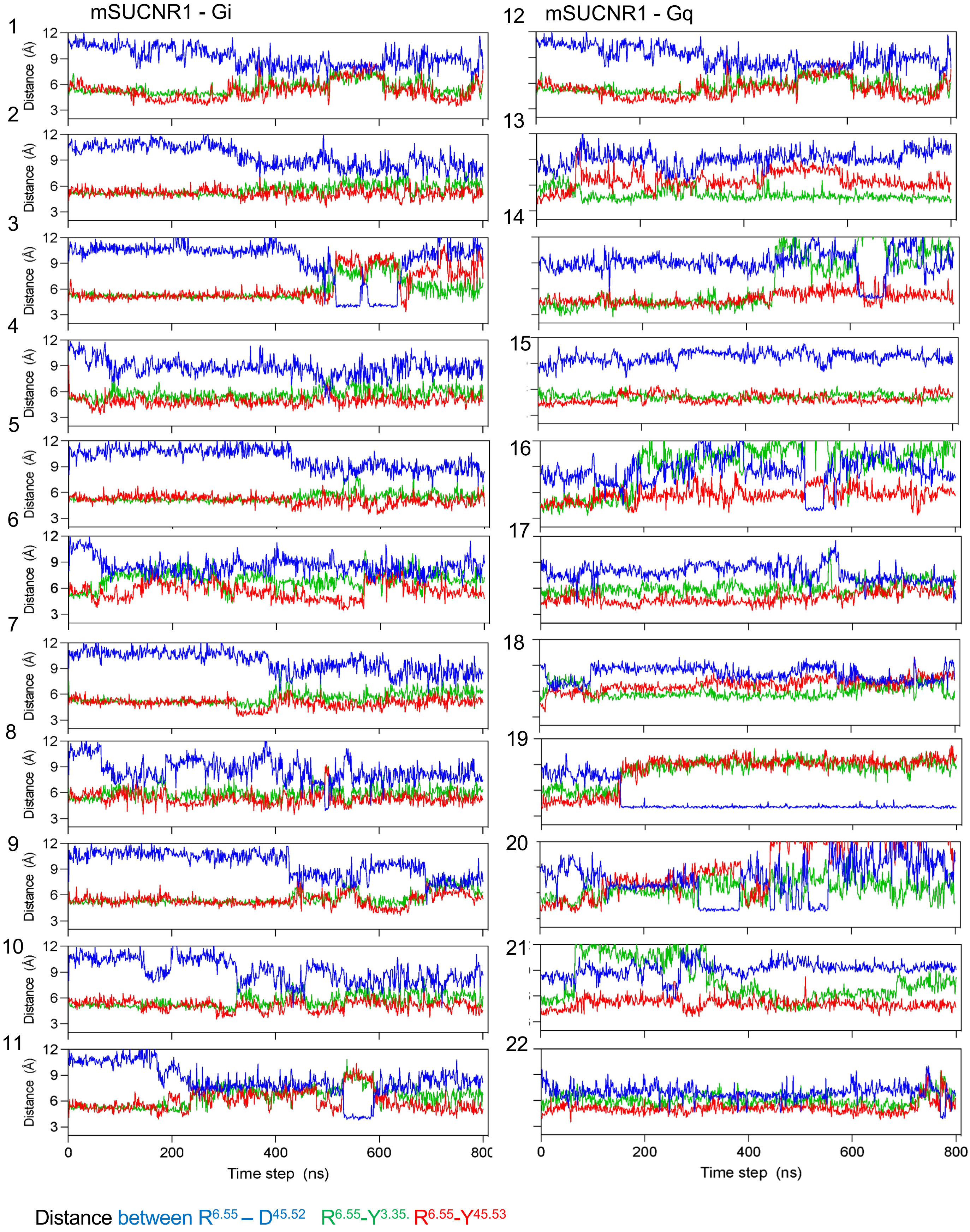


**Supplementary Figure 11.** Distance between $R^{6.55}$ – $D^{45.52}$ (blue), $R^{6.55}$-$Y^{3.35}$ (green), $R^{6.55}$-$Y^{45.53}$ (red) through every simulation **A**. – mSUCNR1 - Gi and **B.** - mSUCNR1 - Gq

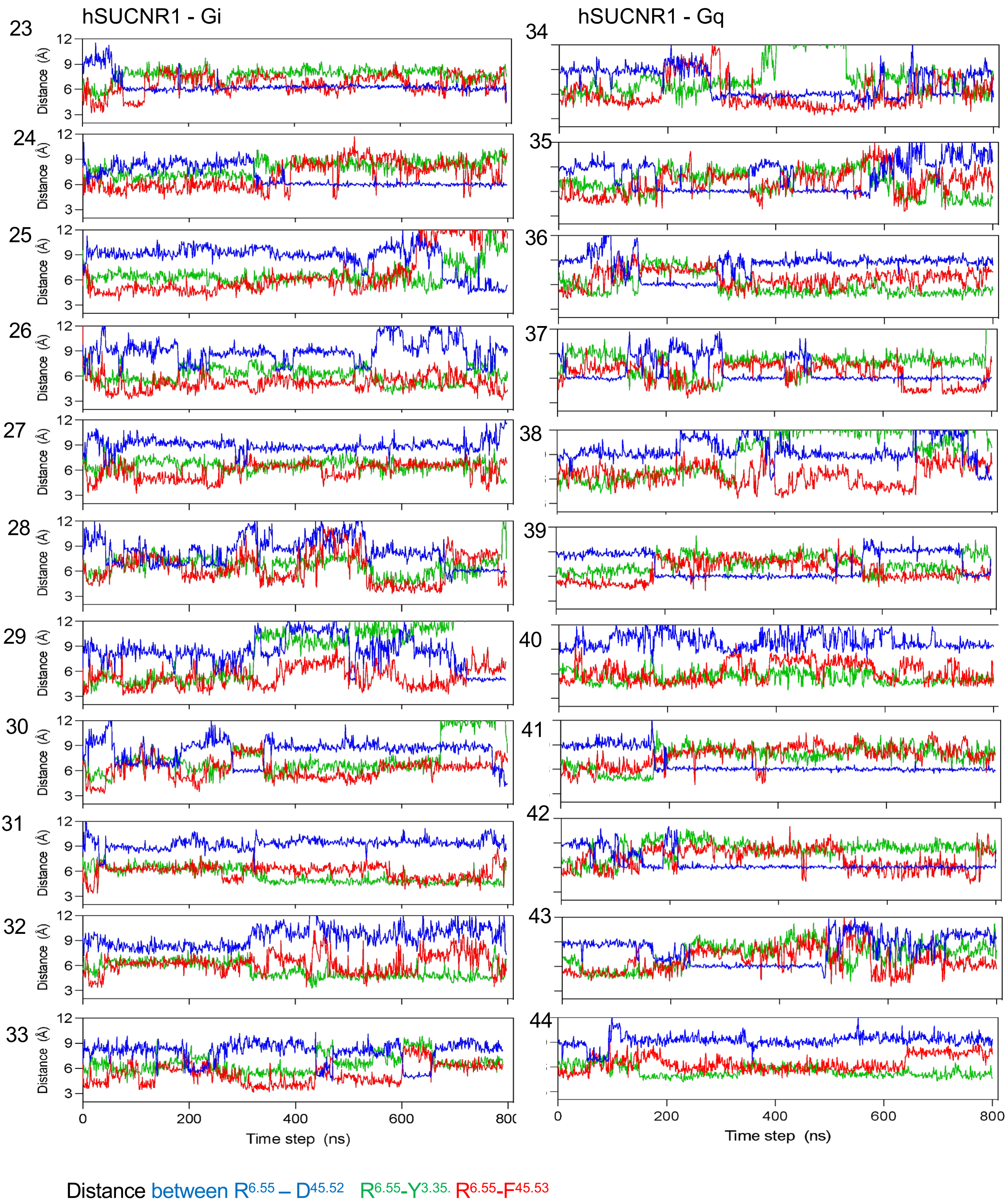


**Supplementary Figure 12.** Distance between $R^{6.55}$ – $D^{45.52}$ (blue), $R^{6.55}$-$Y^{3.35}$ (green), $R^{6.55}$-$F^{45.53}$ (red) through every simulation **A.** – hSUCNR1 - Gi and **B.** - hSUCNR1 - Gq
